# Tuning mechanical *milieux* of tissue templates and their cellular inhabitants to guide mechanoadaptation

**DOI:** 10.1101/2024.12.03.626678

**Authors:** Vina D. L. Putra, Vittorio Sansalone, Kristopher A. Kilian, Melissa L. Knothe Tate

**Affiliations:** School of Chemistry and School of Materials Science & Engineering, University of New South Wales, Sydney, NSW, Australia; Laboratoire Modélisation et Simulation Multi Echelle, Equipe Biomécanique, Faculté des Sciences et Technologie, Université Paris-Est Créteil Val de Marne, France; Blue Mountains World Interdisciplinary Innovation Institute (bmwi³), Blue Mountains, NSW, Australia

**Keywords:** Mechanoadaptation, Mechanomics, Remodeling, Stiffness, **C**ytoskeleton, Development, Computational Fluid Dynamics Modeling

## Abstract

Mechanomics describes the adaptation of mesenchymal stem cells (MSCs) to their mechanical environment, via cytoskeletal remodeling, as well as changes in shape and volume, ultimately resulting in emergent lineage commitment. Here we elucidated effects of exogenous microtubule stabilization, using paclitaxel (PAX), on stem cells’ capacity to sense and adapt to changes in their local mechanical environment. We studied the interplay between the living, evolving cells and their mechanical environment using established experimental and computational tools for respective delivery and prediction of shape and volume changing stresses. Stiffened and volumetrically larger microtubule-stabilized MSCs and their experienced significantly different normal and shear stress compared to control cells when exposed to identical bulk laminar flow (0.2 dyn/cm^2^) for one hour. These spatiotemporal mechanical cues transduced to the nucleus via the cytoskeleton, triggering significantly different changes in gene expression indicative of emergent lineage commitment than those observed in control cells. Using a paired computational model, we further predicted a range of mechanoadaptation responses of microtubule-stabilized cells to scaled up flow magnitudes (1 and 2 dyn/cm^2^). Hence, MSCs adapt to as well as modulate their own mechanical environment via cytoskeletal remodeling and lineage commitment - microtubule stabilization changes not only MSCs’ mechanoadaptive machinery, their capacity to adapt, and their lineage commitment, but also their mechanical environment. Taken as a whole, these studies corroborate our working hypothesis that MSCs and their mechanoadaptive machinery serve as sensors and actuators, intrinsically linked to their lineage potential via mechanoadaptive feedback loops which are sensitive to exogenous modulation via biochemical and biophysical means.

**Classification:** Biological Systems Engineering, Computational Simulations, Cell Biology, Biophysics

## Introduction

The multiscale process of development, from single cells to tissues, to organs, culminating in organisms, is highly mechanobiological^1,2^. Across time and length scales, mechanical cues, along with biochemical cues dictate the emergence of structure and function in developing biological tissues. Single cells proliferate into multicellular constructs and, in the process, undergo transformations involving constrictions, tissue folding, polarization, and condensation, all of which represent hallmarks of specific tissue template creation^3^. The evolution of the multicellular construct exposes the cells within to gradients of biochemical factors and mechanical stresses, presented spatially and temporally, which drive region-specific, tissue level morphogenetic events^4^. As tissue template development progresses, cells within adapt to those cues and thereby modulate the emergent mechanical properties of the tissue template, in a time- and position-dependent manner. Therefore, elucidation of stem cell mechanoadaptation is expected to provide a foundational basis for *mechanomics* engineering of tissues, and to guide tissue neogenesis in a regenerative medicine context.

*Mechanomics* describes the adaptation of stem cells (e.g. MSCs) to their mechanical environment, *e.g.* via cytoskeletal remodeling, as well as changes in cell shape and volume, ultimately resulting in emergent lineage commitment^5^. Physiological stresses experienced by mesenchymal stem cells (MSCs) include coupled volume-and shape-changing stresses, *e.g.* compression and shear. As MSCs adapt their shape, volume and cytoskeleton to these mechanical cues, they also produce extracellular matrix proteins and divide; together these intra- and extracellular adaptations themselves modulate the MSCs’ local mechanical environment, presenting complexities and unique challenges for mechanistic elucidation.

Over the past two decades, a number of tools have been developed to deliver controlled mechanical cues to MSCs and to decipher specific effects of biochemical and mechanical cues in MSC mechanoadaptation^5–16^. Aiming to recapitulate the physiological mechanical environment of MSCs in developing tissue templates, bottom-up approaches have been developed to map cellular responses to specific biochemical and mechanical cues, enabling correlation, in near real time, changes in the stress states and/or mechanical properties of stem cell to their emergent lineage commitment, measured in context of up- and down-regulation of gene transcription.

In previous studies Song et al.^14,17^ developed and validated novel, paired computational and experimental tools for delivery of controlled shear and normal stresses to stem cells. By characterizing the strain response of cells seeded at increasing densities and/or exposed to fluid flow, Song et al. created a platform for mechanical testing of live stem cells as they adapt and as lineage commitment unfold^17^. Increasing seeding density has been shown to induce local compressive stress to MSCs, modulating their nucleus shape, and consequently their fate^8^, while emulating the multicellular proliferation and condensation the during development. Cells seeded at higher densities exhibit flatter nuclei, correlating with increased gene expression for markers of mesenchymal condensation, as well as chondrogenic and osteogenic differentiation^15^. The manner in which density is achieved also profoundly influences MSCs’ nuclear volume and shape, and consequently MSC fate commitment. For example, compared to seeding at target density, stem cells proliferated to target density exhibit rounder nuclei but smaller volume which correlates highly with upregulation of *runx2, sox9* and *aggrecan*, the markers for mesenchymal condensation^8,9,15^. Seeding at increased target density leads to upregulation of pre- and peri-mesenchymal condensation markers, *msx2* and *sox9*, which is cross-correlated with changes in F-actin and microtubule spatial distribution^13^. Together with laminar flow to induce shear and normal stresses on cell surfaces interfacing with the flow field, varying the seeding protocol to alter the rate of achieving target density also influences nuclear shape and modulates the gene expression markers for pre-, peri-, and post-mesenchymal condensation^8,9^. Hence, increasing seeding density provides not only a feasible and tested means to modulate MSC shape and early lineage commitment^9^, but also to emulate the intrinsic processes of emergent tissue template formation and higher length scale tissue genesis for tissue engineering^8,15^.

A number of published studies report effects of flow on the structure and function of endothelial and pluripotent cells for vascular and other tissue engineering applications, *e.g.* changes in shape, cytoskeletal remodelling, and lineage commitment of pluripotent cells to endothelial or vascular-like phenotypes for vascular tissue engineering applications^18^ and to elucidate fibroblast responses to interstitial flow^19,20^. These studies report instantaneous cell responses to flow, as evidenced by 50% reduction in movement of endothelial cells and changes in cell shape within minutes of exposure. After 24 hours’ laminar flow, cells elongated and aligned, accompanied by an increase in F-actin alignment. The rate of actin turnover also increased with an initial decrease in actin polymerization which facilitates F-actin remodelling of individual cells in confluent monolayers exposed to flow^21^. Magnitude of exposure of murine embryonic stem cells (ESCs) to flows ranging from 1.5 to 15 dyn/cm^2^ influences differentiation efficiency towards endothelial and hematopoietic lineages^22^. Given the interdependence of flow-induced cytoskeletal remodeling, cell shape changes and cell differentiation, there is a need to characterize and map emergent changes in multiscale structure and function of live cells as they adapt to flow.

Previous studies demonstrated that the presence as well as the density of cells *per se* influence both local flow regimes as well as stresses experienced by cells at the cell-flow interface^14,17,23^. In other words, cells seeded at various densities influence flow fields in their local milieu which in turn modulates force transduction at interfaces with cells and cell deformation^12,14,17^. Hence, stem cells adapt to as well as modulate their local environment via mechanoadaptation, similar to a mechanical test on live cells where the test conditions guide emergent lineage commitment and nascent tissue template genesis^2^. Developmental processes can be emulated via seeding protocols (seeding versus proliferating to target density), selection of substrate compliance and/or cell adhesion proteins and patterns^16,24–26^, and exogenous delivery of controlled shape and volume changing stresses^14^. Stem cells exhibit significantly greater (1000-fold) sensitivity to mechanical cues than terminally differentiated cells^8,13^; Exposure of MSCs to controlled laminar flow from 0.2 – 2 dyn/cm^27,9,11,13^ elicits significant changes in baseline gene expression of mesenchymal condensation markers indicative of incipient skeletogenesis^8,12,13,15,25,26^.

The cytoskeleton, *i.e.* actin and microtubule polymer filaments, provides both structural scaffolding as well as critical sensing and force generation functions. The unique remodeling of actin and microtubule translates to the evolution of specialized structure and function during MSC differentiation towards e.g. osteogenic, chondrogenic or adipogenic lineages (^1^, reviewed in *Putra et al. 2023*^10^). During exposure to flow, microtubule concentrations per cell change as a function of seeding density and cell height (affecting drag), while changes in F-actin are more influenced by the magnitude and duration of flow exposure^13^. Furthermore, exogenous exposure of stem cells to the microtubule stabilizing agent, Paclitaxel (PAX), induces pronounced structural changes including increased volume and F-actin alignment over time^25,26^, and a corresponding increase in Young’s modulus. These studies, probing effects of exogenous biophysical and biochemical cues on stem cell structure and function and emergent lineage commitment, provided impetus for us to map local stress and strains imbued by flow on cell surfaces and spatiotemporal changes in F-actin and microtubule spatial distribution in the first hour of exposure to such flow, in microtubule stabilized and control cells. Delineating the spatiotemporal emergence of structure and function in developing tissues as a function of controlled exogenous biophysical (volume- and shape-changing stresses) and biochemical cues, is expected not only to refine and expand the mechanome map^14^ but also to provide a reference library of exogenous cues to guide cell lineage commitment and bottom-up genesis of tissues.

Here we aim to probe the role of the microtubule and actin cytoskeleton in stem cell mechanoadaptation to controlled shape- and volume-changing stresses. Our approach was to combine an established MSC model with an established model of microtubule stabilization and a previously developed system for controlled flow and delivery of shape (shear) and volume-changing (normal) stresses to cells seeded within to test the hypotheses that:

- microtubule-stabilized cells will experience different magnitudes and spatial distributions of normal and shear stresses compared to control cells exposed to identical flow regimes, due to associated changes in volume and stiffness of microtubule-stabilized cells,
- microtubule-stabilized cells seeded at increasing seeding density will both experience different effects of flow and will themselves influence their local mechanical milieu differently, where at higher seeding densities, cells may experience smaller normal and shear strains,
- these differences in normal and shear stress magnitude will be transduced to the nucleus via the cytoskeleton, resulting in differences in lineage specific gene expression.

In addition, we used a previously tested and validated computational model of the experimental set up to expand our hypothesis testing to a range of variables, predicting effects of decreased microtubule stabilizing agent (PAX concentrations at 1 nM, 10 nM) and increased flow magnitudes (1 and 2 dyn/cm^2^). Insodoing, we aim to further elucidate the stem cell mechanome, further developing stress-strain-fate maps to build a foundation for prospective delivery of mechanical cues to achieve targeted cell fates and emergent tissue types.

## Materials and Methods

### Experimental Methods

#### CHOICE OF MESENCHYMAL STEM CELL MODEL

To leverage our previous libraries of published mechanome data, we used the model murine embryonic mesenchymal stem cell line (C3H10T1/2 cell line, ATCC, CCL-226) for this series of experiments. This conferred important advantages with regard to both validation as well as comparability with previously published data sets from several series of stem cell mechanomics studies^12,14,17^

#### CELL CULTURE

Cells were maintained in culture and used in experiments between passages 5 and 15, according to our previous protocols^13^. Cells were passaged in standard culture medium (Basal media eagle supplemented with 10% foetal bovine serum (FBS), 1% L-glutamine, and 1% penicillin/streptomycin (Invitrogen, Carlsbad, CA)) in T75 culture flasks (Corning Inc. Corning, NY). These cells were detached using 0.25% Trypsin-EDTA (Invitrogen, Carlsbad, CA) for 5 minutes, centrifuged, resuspended in fresh culture medium, and seeded on glass coverslips of 25 mm diameter. To exclude biochemical and/or other potential inductive effects on adhesion or other cell behavior, no extracellular or other matrix coating was used.

#### INDEPENDENT VARIABLES: TARGET SEEDING DENSITY AND MICROTUBULE STABILIZATION

Cells were seeded at specific increasing seeding densities shown previously to induce local compression on cells seeded at target density, i.e. low density (LD 5,000 cells/cm^2^), high density (HD 15,000 cells/cm^2^) or very high density (VHD 45,000 cells/cm^2^). A subset of cells were exposed to an exogenous microtubule stabilizing agent, paclitaxel (PAX, molecular weight = 853.906 g/mol). According to previously established protocols^25,26^, a 1.2 μl of PAX stock of 1 mg/ml in dimethyl sulfoxide (DMSO) was added to 14 ml of medium to make 100 nM PAX solution. 24h after cell seeding, the medium was replaced with PAX containing medium and incubated for 72h.

#### FLOW EXPERIMENT

Pre-warmed (37°C) phenol red-free and serum-free culture medium was used as flow medium to exclude potential interaction effects of FBS and flow, as per previous published protocols^12,14,17^. The viscosity of serum-free culture medium is close to that of physiological saline (0.001 kg/ms). The medium was pumped through the ProFlow chamber (Warner Instruments) using a 10 ml syringe (Terumo) in a screw pump system (Harvard Apparatus, Holliston, MA).

The ProFlow chamber was designed, tested and validated to expose cells to controlled flow-induced shear and normal stresses at specific magnitude and durations, to mimic the effect of shape changing, deviatoric stresses and volume changing dilatational stresses, on the stem cells, analogous to previous protocols^8,13,14^. For this set of experiments, a target flow magnitude of 0.2 dyn/cm^2^ was chosen as previous studies showed this to be the smallest predicted stress magnitude that could induce significant changes, compared to baseline, in gene expression of MSCs^2^. Cells seeded on a 25 mm diameter coverslip were fixed in the bottom plate of the chamber and covered with silicon gasket with a 15 mm diameter coverslip fixed on the top plate. The tops and bottom plates were assembled and tightened with 4 thumb screws. The flow rate was set at 0.134 ml/minute to deliver 0.2 dyn/cm^2^ as previously predicted and described^13,14,17^. Cells cultured analogously but not exposed to flow (static) served as a baseline controls.

#### LIVE IMAGING OF CYTOSKELETON AND CELL DISPLACEMENTS UNDERFLOW: For tracking live

cell displacement of cell surfaces under flow (digital image correlation, DIC), cells were labelled with 1 μm diameter microbeads (Bangs Lab) adsorbed with concanavalin-A (con-A) and conjugated with flash red fluorophore 620 nm according to previously published protocols^14^. 50 μl of bead solution (10 mg/ml beads-con-A stock) was added to the cells per well (6-well plate), in serum-free medium, and incubated for 30-40 minutes. The coverslip with seeded cells was then transferred to the flow chamber for further live imaging.

For live cytoskeleton imaging, cells were labelled with actin SPY-620 nm probe and microtubule SPY-555 probe (Cytoskeleton Inc.) by adding 2μl probe solution per ml medium for 30 minutes prior to transferring the coverslip into the flow chamber for flow exposure. Using the Zeiss Elyra 7 Lattice SIM microscope, cells were imaged in 4D (xyzt) at t = 0 (just prior to flow exposure), and subsequently at t = 300 seconds, and for every 5 minutes during the 60 minutes’ flow exposure. Hoechst labelled nuclei, actin, and microtubule were imaged using 405 nm, 650 nm and 561 nm diode laser respectively. Laser power was set at 50 mW (milliwatts). A 40x, 1.3 NA oil immersion objective lens was used, resulting in xy image size of 126.21 µm x 126.21 µm. 4D images were acquired by selecting z-stack mode with leaping step size of 0.273 µm.

#### RT-PCR (REVERSE TRANSCRIPTION POLYMERASE CHAIN REACTION)

Relative expression of gene markers indicative of early mesenchymal condensation was probed using rt-PCR, analogous to our group’s previously published approach^9,13,14^. Total RNA was extracted from cultured cells exposed to 0.2 dyn/cm^2^ flow regime within the flow chamber, using an RNA extraction kit (Life Tech) according to the manufacturer’s instructions. Cells from 5-6 coverslips/chamber were pooled to generate sufficient RNA. 500 – 1000 ng of total RNA isolated per 20 μl of reaction volume was reverse transcribed into cDNA using the SuperScript First-Strand Synthesis System (Invitrogen). Real-time polymerase chain reactions (PCRs) were performed and monitored using the SYBR Green PCR Mastermix and the 12K-Flex QuantStudio detection system (Applied Biosystem). cDNA samples (1 μl of 10 ng/ul for total volume of 25 μl per reaction) were analysed for genes of interest and the expression was normalized using the reference of TATA binding protein (*tbp*) (**Table 1**). The level of expression of each target gene was then calculated as −2^ΔΔCt^ as previously described^13^. Each sample was repeated at least three times for each gene of interest. rt-PCR was performed at 95°C for 2 minutes followed by 34 cycles of 30-second denaturation at 95°C, 30 seconds of annealing at the primer-specific temperature, and 1 minute of elongation at 72°C.

**Table 1.**
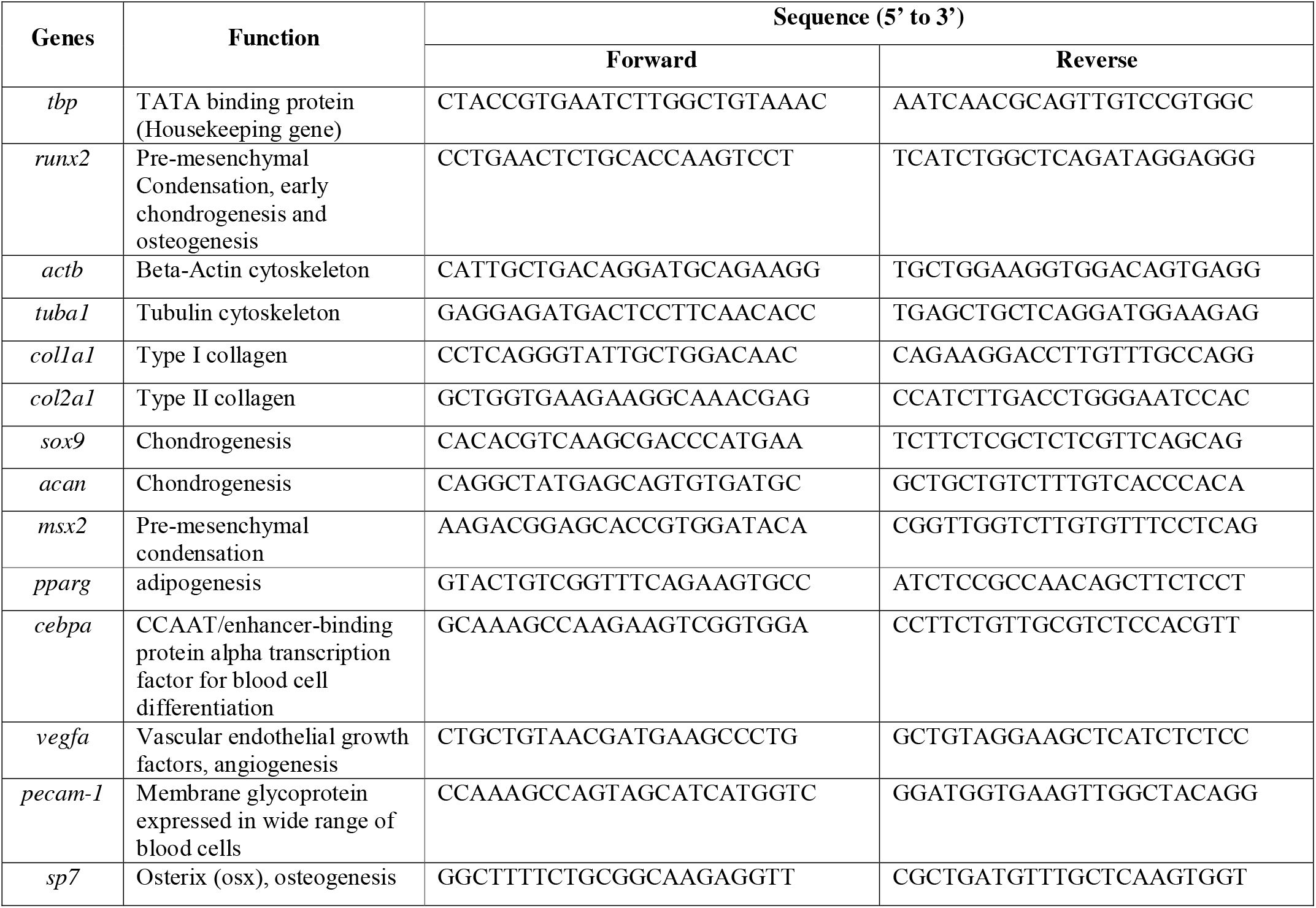
Sequences of primers used in. RT-PCR

#### ANALYSIS OF STEM CELL MECHANICAL PROPERTIES AND STRESS-STRAIN MAPPING

Image analysis was performed using the spot detection function of the software Imaris (Oxford Instruments), which allows tracking of the velocity, position and displacement in x, y and z of individual microbeads and throughout the time series of imaging. The position data were analysed to create a 4D plot in MATLAB to depict the magnitude of displacement of the microbeads over time. The position data at time 0 (min, t_0_) was used as a reference to calculate the original position of the microbeads from which the bead displacement was calculated at each time point. Using idealized linear elastic theory, the elastic modulus data from AFM measurements were used to calculate the stress experienced by the cells at various timepoints during flow exposure.

#### STATISTICAL ANALYSIS

A multivariate correlation analysis was conducted to test interacting effects of probed gene marker expression as well as the expression level and spatial distribution of F-actin and microtubule cytoskeleton (SPSS, IBM). Significant differences in the fold change of gene expression were analysed using two-way ANOVA with Tukey’s multiple comparison test, where a significance level was set at p < 0.05. Multivariate analysis and test between subject effects were performed to analyse the level of interactions between all independent variables in this study: PAX treatment, seeding densities, and flow, in influencing the up- or downregulation of specific genes.

### Computational Methods

Computational modeling of cells and flow within the flow chamber was then used to further elucidate the experimental data and to probe further hypotheses, e.g. regarding effects of higher magnitude flows. Model parameters and predictions were chosen to match those of previously published work, for validation and further expansion upon the previously published, paired computational-experimental studies from our group^13,14,17^.

3D CELL MODEL GEOMETRY DEFINITION AND MATERIAL ASSIGNMENT: Images from the live confocal imaging were processed and a binary mask was applied to identify and isolate the cell bodies. The smoothed 3D cell objects were exported from Simpleware ScanIP (Version 2018.12, Synopsys, Inc., Mountain View, USA) as STL files and then imported into Ansys® ICEM CFD 19.2 for meshing (Fig. S5.4A). This smoothing process requires elimination of small protrusions or irregular complex, structures (less than 0.5 μm) near the cell body that may interfere with the assignment of materials and physics. The mesh models were created with refined Prism meshes around the cell-fluid interfaces so that the computational efficiency was ensured without compromising simulation accuracy. The final average degree of freedom (DoFs) was about 1,654,313 with 442,812 domain elements assigned to cells and peri-cell regions.

Meshed cell models were imported into COMSOL as a mesh for computational fluid dynamics analysis of normal and shear stresses at cell-flow interfaces. A sub-model of the microenvironment of the cells within the chamber was created including the mesh of the cell structures and surrounding medium, within a 400 x 400 x 250 μm bounding box. Meshes were imported and material specifications (Supplementary table 5.1) assigned to parts of the mesh consisting of fluid (cell culture medium) with density and viscosity similar to that of saline, the glass boundary at the top and bottom of chamber, and the cells seeded therein. All computational modeling methods conformed to those published and validated previously.^11,17^

### MEDIUM FLOW MODELING AND SUB-MODELING OF CELLS WITHIN CHAMBER: COMSOL

Multiphysics software was used in the modelling of the flow chamber and computational fluid dynamics analysis to deliver computationally simulated flow and to predict the flow fields around the stem cells in the Proflow chamber designed^11^ to deliver highly controlled shear stress in the middle 80% area of the chamber^11,14^. The chamber is an eye-shaped structure based on the geometry of the gasket with the thickness of 250 μm, 23 mm length and 11 mm width, the inlet and outlet located in both corners of the chamber. A no-slip wall condition is assigned at the walls of the chamber. For all models, the continuity equation (1) and Navier-Stokes equations (2) are solved using a 2^nd^ order upwind-discretization scheme in three dimensions. The Navier Stokes equation, 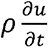 and *u*. ∇*u* represent acceleration. [- ∇pI + K] represents the viscous force for incompressible fluid, and F represents the external force, or external forces applied to the fluid^27^. The wall shear stress is calculated from the wall strain rate (3) based on the flow between two parallel plates, as the wall shear stress relates to velocity by viscosity (μ) and the rate of strain (the rate of change in velocity over the distance across y-axis) (Fig. S5.4B). Flow is assumed to be laminar, incompressible, and steady in three dimensions, and the fluid (cell culture medium or saline) was idealized as Newtonian with 996 kg/m^3^ density, 0.001 kg/ms viscosity, 301 K body temperature, according to previous protocols by Anderson et al. 2006^11^ and Song et al. 2012^14^.

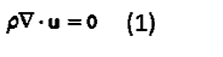

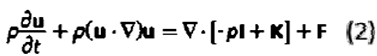

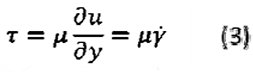

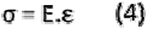

Where u is velocity, ρ is density, μ is viscosity, and τ is shear stress at or is the strain rate and y is the distance from bottom of chamber. To calculate cell stresses and strain, cells were idealized to follow linear elastic theory (4), where E = elastic modulus, σ and ε indicate stress and strain. The linear elastic idealization of the cell mechanics can be adjusted in COMSOL to better capture plastic and viscous behavior of the cells, as addressed in later permutations of the model.

As a first approximation, the elastic moduli for PAX-treated and nontreated cells were assigned based on atomic force microscopy (AFM) measurements of cell stiffness^25,26^. In the previous study by Song et al 2012^14^ parametric estimation of cell elastic moduli was performed by predicting the behaviour of cells with elastic moduli ranging in different orders of magnitude and comparing the computationally predicted stress-strain data with the measured data^14^. In this study, specific elastic moduli values from the AFM measurement were used to define the cell material properties. Steady state, laminar flow was modelled using a Reynolds number of 16 (Reynolds number is the ratio of fluid inertial forces and viscous forces which in this case indicates flow is likely to be laminar). There are numerous other factors that could affect a cell change in volume, such as Poisson ratio (Table S1), which indicate how cells are able to resist compression or volume change under a given tension or compression. As a biological material, cell exhibit a complex material property that behaves as solid or fluid depending on their environment. Consistently, the actin cytoskeleton has been shown to exhibit such adaptive rheology, tuning its fluid-like or solid-like properties, in response to substrate stiffness. In this study, we idealized the cell as having a constant Poisson ratio of 0.36 based on the AFM measurement of cells on glass.

A parametric sweep of independent variables was performed, while introducing an input pressure gradient at the inlet in the range of 4, 20, and 40 Pa, the pressure predicted to achieve the target shear stress (τ) of 0.2, 1, and 2 dyn/cm^2^ within the chamber, which was shown in previous studies to elicit significant changes in gene expression in MSCs^9,13^. Sub-modeling of the flow chamber was done to capture the middle-most area of the chamber where flow velocity and shear stress are most uniformly distributed. This 400 x 400 μm area with the chamber height of 250 μm was modelled as a box containing the cell structure adherent to the inner bottom coverslip of the chamber (Fig. S5.4C, D, E). The flow specification (detailed in Supplementary Table S2) within this sub-model was assigned as fully developed laminar flow with velocity of 0.00011 m/s as modelled in the full flow chamber at the location x = 11.5 mm, y = 5 mm, and z= 0.01 mm per previous model specifications^11,14^.

The velocity and shear stress at the vicinity of the single cell surface are calculated by selecting a coordinate in the ‘Cut Point 3D’ function (Fig. S5.4F). The velocity profile at the apical-basal height of the cells seeded at higher densities is calculated by selecting the z-slice in the 3D model at 0, 2, 4, 6, 8, and 10 μm, then computing the velocity (*spf.U*) using the ‘max/min surface function’ (Fig. S5.4G). At increasing height of the chamber, the shear rate is defined as the gradient in velocity, that is, the difference in velocity between the two surfaces containing the fluid, divided by the distance between them.

## Results

### Effect of microtubule stabilization and flow exposure on cells

C3H/10T1/2 cells treated with 100 nM PAX were previously shown to exhibit pronounced structural changes involving volume increase and F-actin alignment while being under the challenge of proliferation due to stabilized microtubule^25,26^. Hence, we hypothesized that microtubule-stabilized cells will experience different magnitudes and spatial distributions of normal and shear stresses compared to control cells exposed to identical flow regimes, due to associated changes in volume and stiffness of microtubule-stabilized cells. Further, we hypothesized that these differences in normal and shear stress magnitude will be transduced to the nucleus via the cytoskeleton, resulting in differences in lineage specific gene expression (addressed below).

Control and microtubule-stabilized cells were exposed to flow at 0.2 dyn/cm^2^ using the Proflow chamber (Fig. 1A) to allow the study of their structural adaptation under the flow induced shear stress. Spatiotemporal maps of flow-induced strain fields were generated by tracking displacement magnitudes and directions for fluorescent microbeads tagged to cell surfaces and calculating strain in the xy and xz planes (Supplementary video S5.1). We observed both immediate passive effects of flow as well as adaptation of cell shape and height within 45 minutes of flow exposure (Fig. 1B). The positions of each bead were tracked and mapped for their magnitude and direction over time (Fig. 1C).

**Figure 1.**
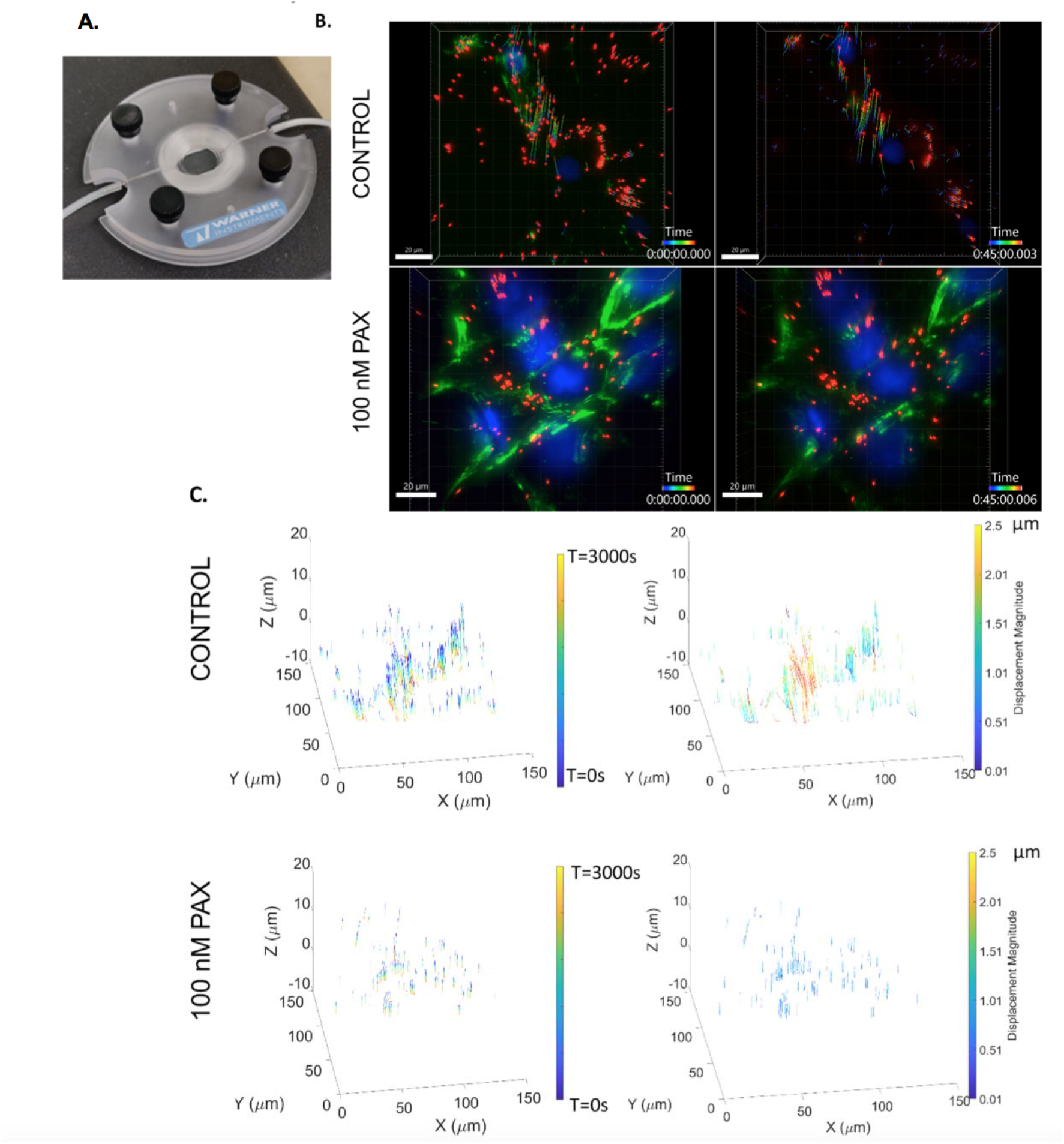
Measurement of bead vector displacements in three dimensions (xyz) and over time during exposure to controlled magnitude (0.2 dyn/cm^2^), laminar flow. (A) Experimental set up with flow chamber designed and tested to deliver controlled laminar flow to cells^14,17^. (B) Control and cells exposed to 100 nM PAX, at time zero (left) and after 45 minutes’ flow exposure (right). After 45 minutes of flow, control cells exhibited larger bead displacements and over a more extended period of time of surface attached beads as cells were adapting closer to the substrates, compared to those of PAX treated cells. (C) (left) Bead displacement (xyz) over time (seconds). (right) Bead displacement magnitude (xyz) in µm.

### Microbead displacements in response to flow and effects of microtubule stabilization

Flow-induced microbead displacements were larger in control cells without microtubule stabilization compared to microtubule-stabilized cells. In response to flow, beads attached to control cells gradually moved toward the substrate over time. Microtubule-stabilized cells showed smaller magnitude microbead displacements in the z direction. This study reports on stem cell adaptation to controlled, steady state, laminar flow for cells seeded at different target cell seeding densities and effects of exogenous microtubule stabilization using PAX. Hence, time zero captures the xyz position of cells and surface attached microbeads at the earliest observation point after initiation of flow.

All xyz measurements of displacements were normalized to those of the static (non-flowed) condition (Supplementary Video S5.2), where microbead displacement remain constant over time (Fig. S1A). The microtubule-stabilized static group gradually achieved minimal positive z displacement (Fig. S1B). As an additional null control, cells were fixed chemically and exposed to flow and were exposed to flow (Fig. S1C, D). Microbead displacements were minimal in chemically fixed cells.

### Surface strain on flow-exposed cells significantly influenced by both microtubule stabilization and seeding density

Increasing seeding density was previously shown to induce local compression, emulating the multicellular proliferation and condensation the during development^8,9^. Previously, it was also shown that cells seeded at high and very high density exhibit higher stiffness or bulk modulus that was enhanced with microtubule stabilization. Hence, we hypothesized that cells seeded at increasing seeding density will both experience different effects of flow and will themselves influence their local mechanical milieu differently, where at higher seeding densities, cells may experience smaller normal and shear strains.

Cells exposed to flow experience both normal and shear stress, and therefore we calculated those stresses component based on the microbeads displacements in XY (accounting for stress parallel to the direction of flow) and XYZ (accounting for tensile and compression elements).

Both microtubule stabilization as well as seeding density had profound, complex and interacting effects (see correlation analysis below) on surface shear and normal strains experienced by stem cells exposed to controlled laminar flow. The increase in cell volume associated with microtubule stabilization appeared to attenuate shear strains on cells (Fig. 2A) compared to control cells. At (low density) LD, flow-induced bead displacement occurred in multiple directions within the xy-plane. At higher seeding density (HD), both control and PAX-treated cells exhibited a larger and more random displacements in the xy-plane (Fig. 2B). This was likely due to the effect of boundary “smoothing” at the fluid-cell interface of larger cells compared to that of an equivalent surface of multiple smaller control cells. Similarly, the increase in cell stiffness associated with microtubule stabilization appeared to enable the cell to better resist flow-induced strains, as observed by decreased normal strains at cell surfaces (Fig. 2C) (Supplementary Video S3). Cell height (measured as mean z-position of microbeads) decreased in cells exposed to flow compared to the static condition (Fig. S2A). Microtubule stabilization exerted a significant effect on cell height during flow exposure, compared to non-treated control cells (Fig. S2B, C, D).

**Figure 2.**
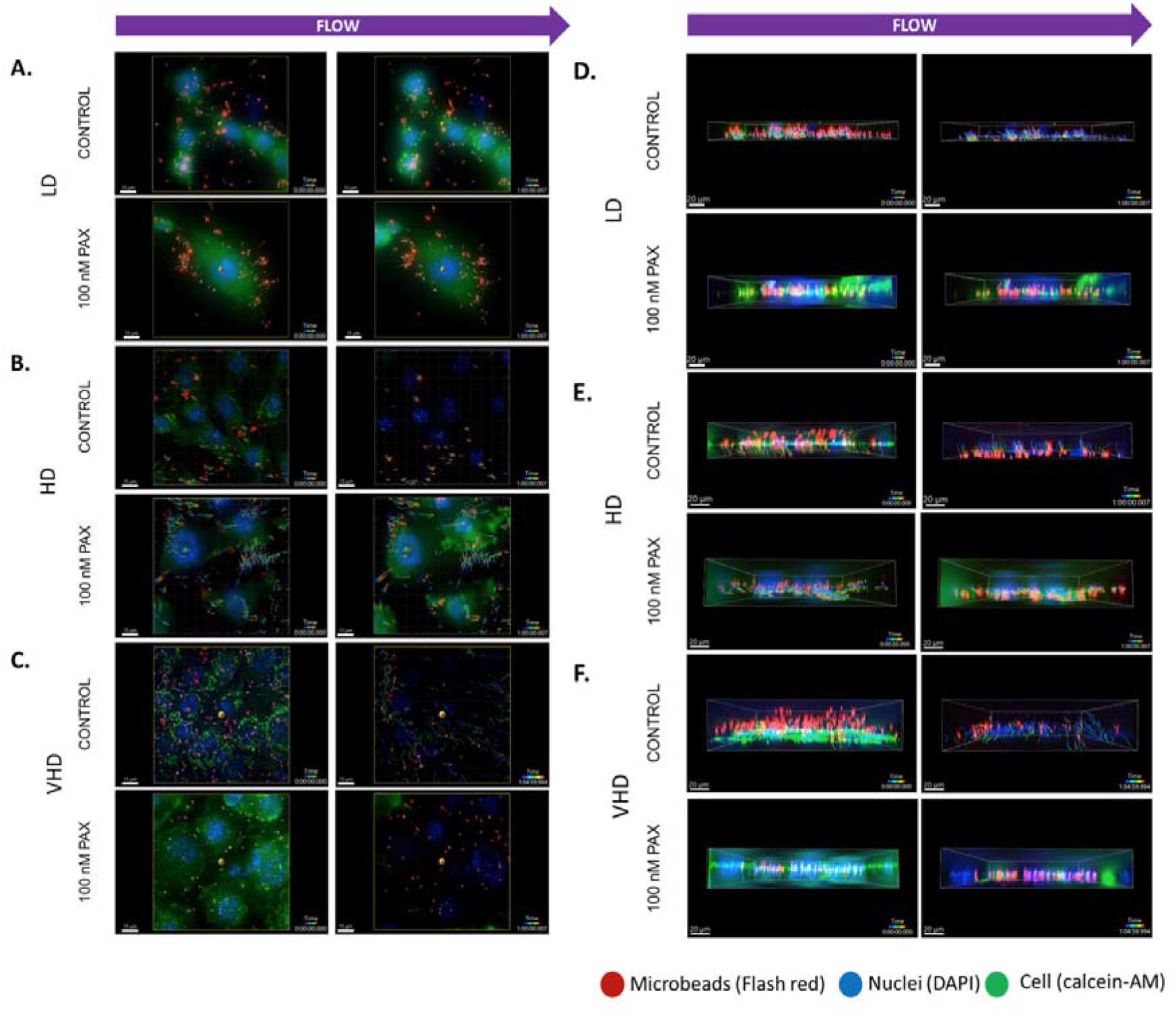
Effect of seeding density and microtubule stabilization (PAX) on bead displacements under flow ^14^. Cells seeded at increased density were exposed to flow (arrow indicates flow direction from left to right). Control and microtubule-stabilized stem cells (100 nM PAX) seeded at low density (LD, 5000 cells/cm^2^) (A), high density (HD, 15000 cells/cm^2^) (B), and very high density (VHD, 45000 cells/cm^2^) (C) exhibit different degrees of adaptation to flow at the subcellular scale (scale bar = 15 μm), where increasing cell-surface-attached-bead displacement with time is indicative of cell adaptation. Bead displacement in the xz plane, observed in micrographs of cells seeded at LD (D), HD (E), and VHD (F) and exposed to flow over time.

**Figure 3.**
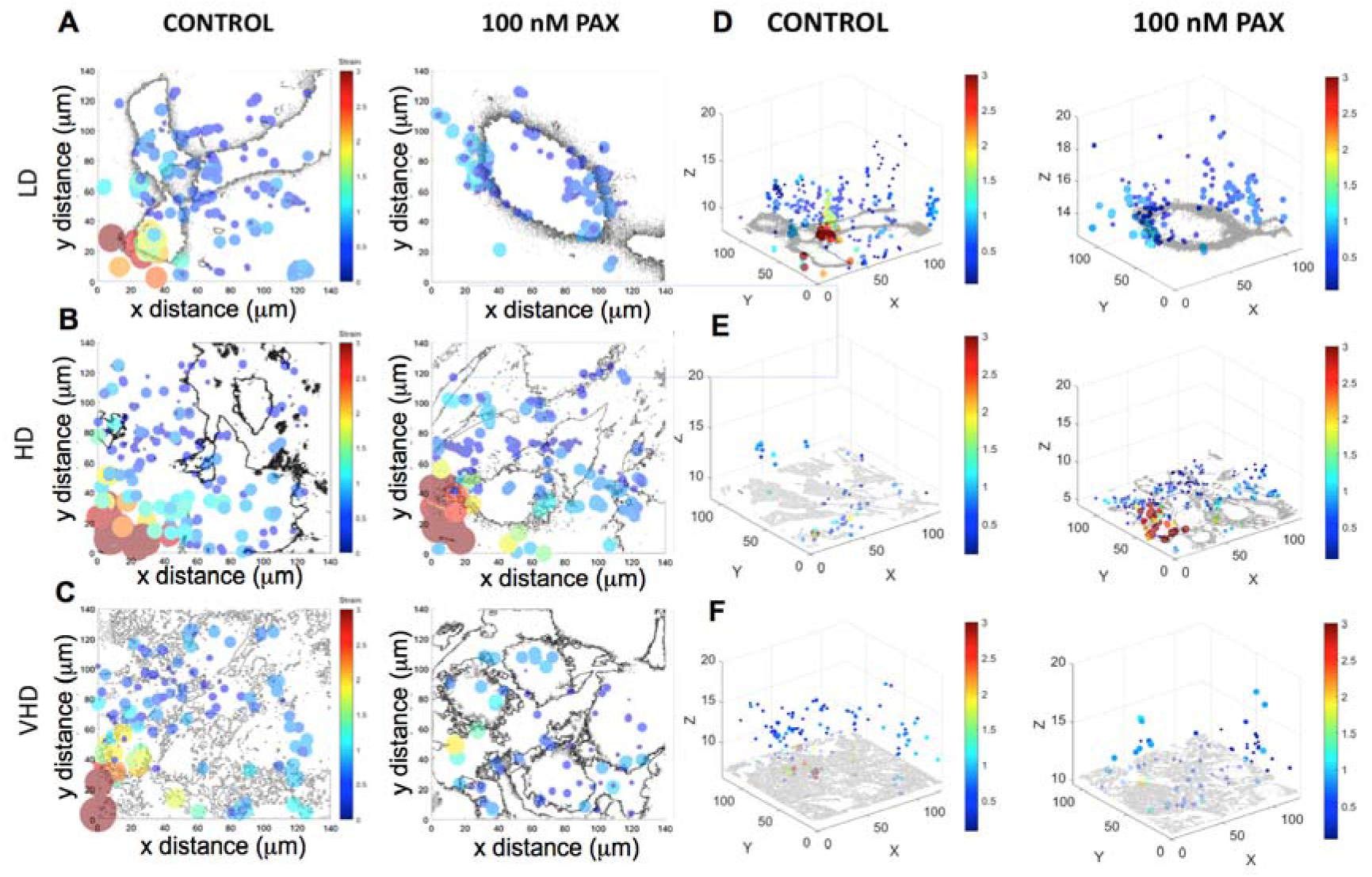
Effect of seeding density and microtubule stabilization (PAX) on flow-induced shear and normal strain at cell surfaces ^14^ (experimental setup as shown in Figure 2). Shear strains on surfaces of cells seeded at low density (LD, A), high density (HD, B), and very high density (VHD, C) were calculated based on bead displacements in the xy plane over the 60-minute observation period. Normal strains on surfaces of cells seeded at LD (D), HD (E), and VHD (F) were calculated based on bead displacements in the xz plane over the 60-minute observation period.

Flow-induced displacements in the z-direction were in general larger for all groups than displacements in xy plane (the direction of flow, Fig. S2.E and Fig. 5.2D-F).

The xy bead position of the beads were used to generate the shear strain maps using DIC (Fig. S2F-H). Control cells exhibited larger xy strains than the microtubule-stabilized cells for all LD (Fig. 3A), HD (Fig. 3B), and VHD (Fig. 3C) groups, and these strains were larger toward the flow (front) than away from the flow (back) side. The xy-z position of the beads was then used to generate normal strain maps using DIC. LD (Fig. 3D), HD (Fig. 3E), and VHD (Fig. 3F) cells experienced various degrees of normal strains that also depend on changes in cell geometry due to microtubule stabilization and proximity to the flow front.

In general, within the multicellular fields of view, surface strains closer to the flow inlet and cellular boundaries, exhibited higher local strains, with up to 3% strains observed locally in areas closest to the inlet and cell boundaries. Aside from those local areas of “strain concentration”, strains were generally observed below 1.25%, across groups.

### Microtubule stabilization increases actin and microtubule expression and remodeling

We then hypothesized that the actin and tubulin cytoskeleton will remodel in response to flow exposure, and in interaction with microtubule stabilization and cell seeding density.

Microtubule-stabilized (PAX) cells showed a significant increase in *actb* and *tuba1* gene expression, whereby the increase in *tuba1* was higher than that of *actb* (Fig. 4A). Exposure of microtubule-stabilized cells to flow was associated with a decrease in *tuba1* expression in LD seeded cells which was nearly abrogated in cells seeded at HD and VHD. In contrast, exposure of microtubule-stabilized cells to flow was associated with an increase in *actb* expression in LD seeded cells which was nearly abrogated in cells seeded at HD and VHD.

**Figure 4.**
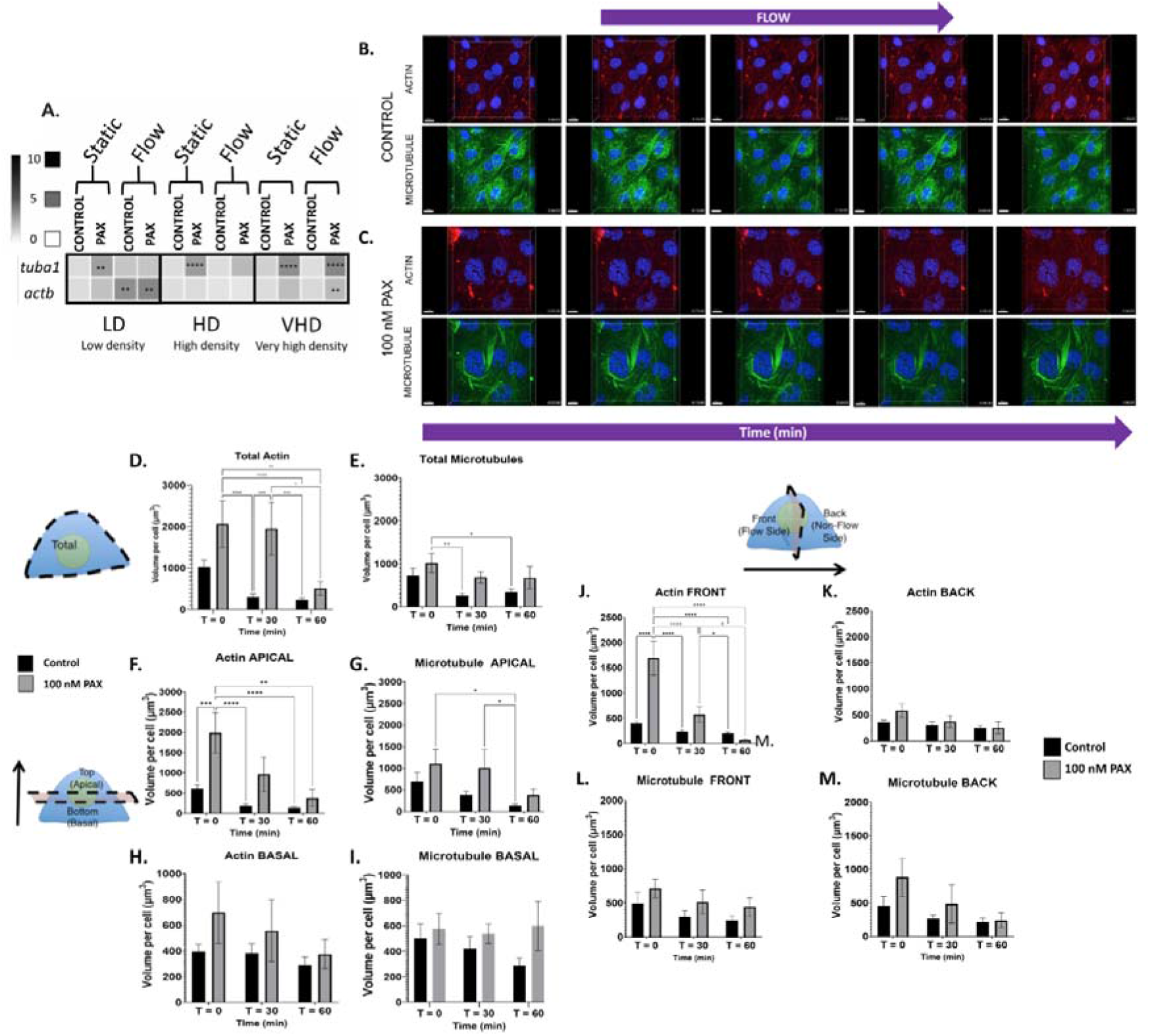
Effect of microtubule stabilization and exposure to flow on actin and microtubule expression and remodeling. Actin and microtubule expression and remodeling, as measured at the mRNA and protein level. (A) Microtubule-stabilized cells exhibit upregulation of both actin (actb) and tubulin (tuba1) mRNA at low density (LD), by 2- and 3-fold respectively, which is nearly abrogated with increasing seeding density. Exposure to flow alone resulted in a significant increase in actb at LD by 3-fold, diminishing to nil at HD. Microtubule-stabilized cells exposed to flow also allows the increase of actb at LD cells by 3.5-fold, but reduced at VHD by 2-fold. Stabilization of microtubule with PAX consistently led to upregulation of tuba1, but not actb, across all seeding density groups. The upregulation of tuba1 was higher in VHD under flow (3.5-fold). Fold change in mRNA was normalized to those of control static group. Asterisk(s) indicate significant up- or downregulation when normalized to the static groups at *p<0.05, **p<0.01, ***p<0.001, and ****p<0.0001 with one-way ANOVA. (A) Micrographs depicting mechanoadaptation via remodeling of F-actin and microtubule filaments in response to flow.

**Figure 5.**
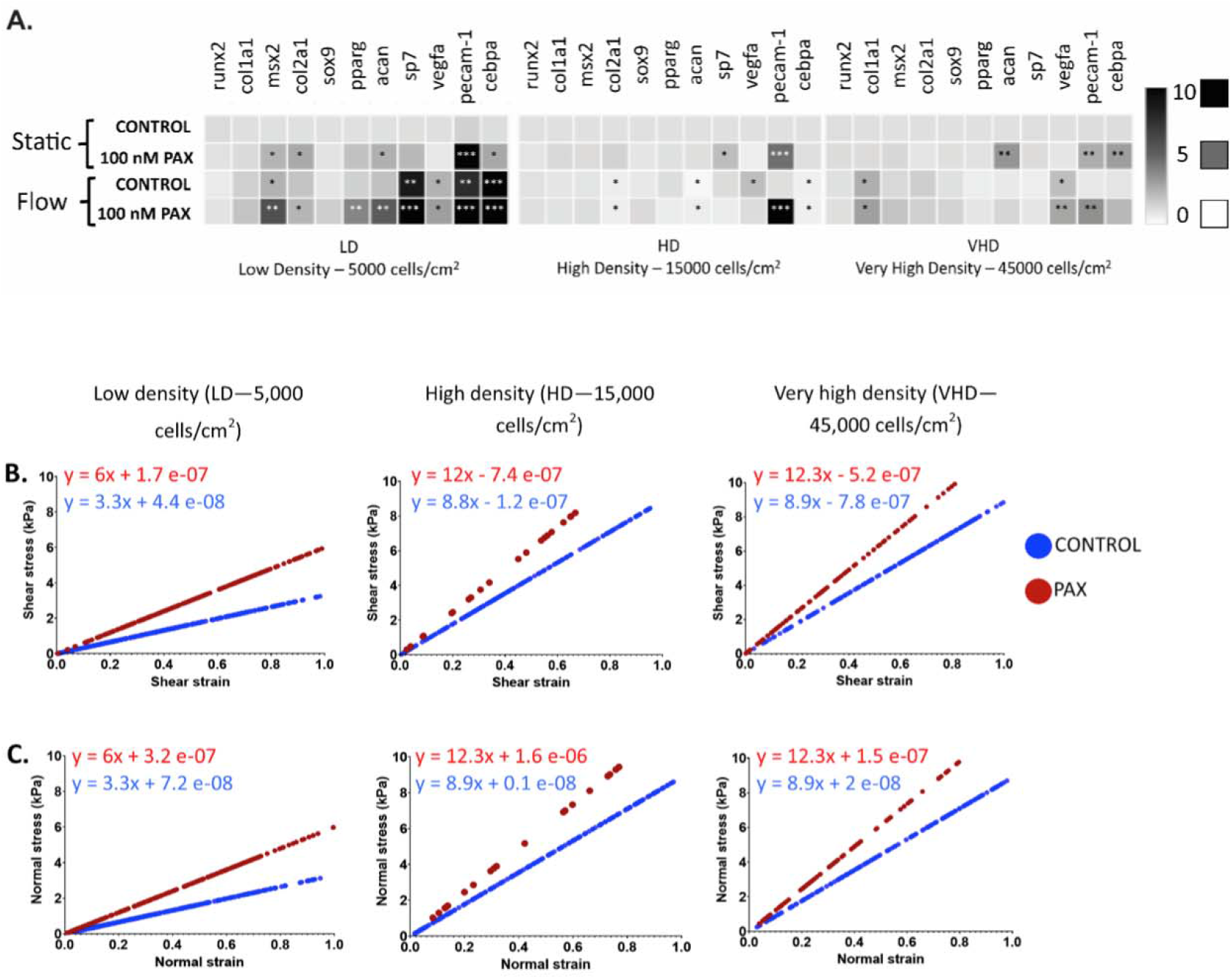
Mechanomics mapping of C3H/10T1/2 murine embryonic stem cells seeded at low density (LD), high density (HD) and very high density (VHD). Cells were exposed to 0.2 dyn/cm^2^ laminar flow. A gene expression profile of emergent changes is linked to stress-strain curves of live stem cells during early fate emergence. (A) Compared to naive control cells seeded at LD, exposure of microtubule-stabilized stem cells to flow significantly upregulated by 2-fold the msx2, col1a1, and col2a1, which are the markers for pre-, peri and post- mesenchymal condensation markers respectively. Microtubule stabilization led to 2-fold upregulation in the osteogenic marker, sp7, in both LD and HD cells, and 3-fold upregulation in the chondrogenic marker, acan, in VHD cells. Under flow exposure, sp7, was also upregulated up to 8-fold and 14-fold in LD control and microtubule-stabilized cells (PAX). However, in microtubule-stabilized and flowed cells seeded at HD, col1a1, sp7 and acan were downregulated by 2-fold. Exogenous stabilization of microtubules (PAX) alone upregulated pecam-1, however, the increase became less significant with increasing seeding densities, HD and VHD. Under flow, pecam-1 and cebpa were upregulated in LD cells by 23- and 20-fold respectively, however, at higher seeding densities, this increase was reduced. Microtubule-stabilized cells exposed to flow consistently upregulated vegfa in all seeding density groups. Asterisk(s) indicate significant up- or downregulation when normalized to the static groups at *p<0.05, **p<0.01, ***p<0.001 with one way ANOVA. In the mouse, mesenchymal condensation occurs at E11.5 or after 11.5 days gestation^2^. (B) Shear and (C) normal stress and strain distributions of stem cells under flow depend on cell seeding density and exogenous microtubule stabilization (PAX).

**Figure 5.**
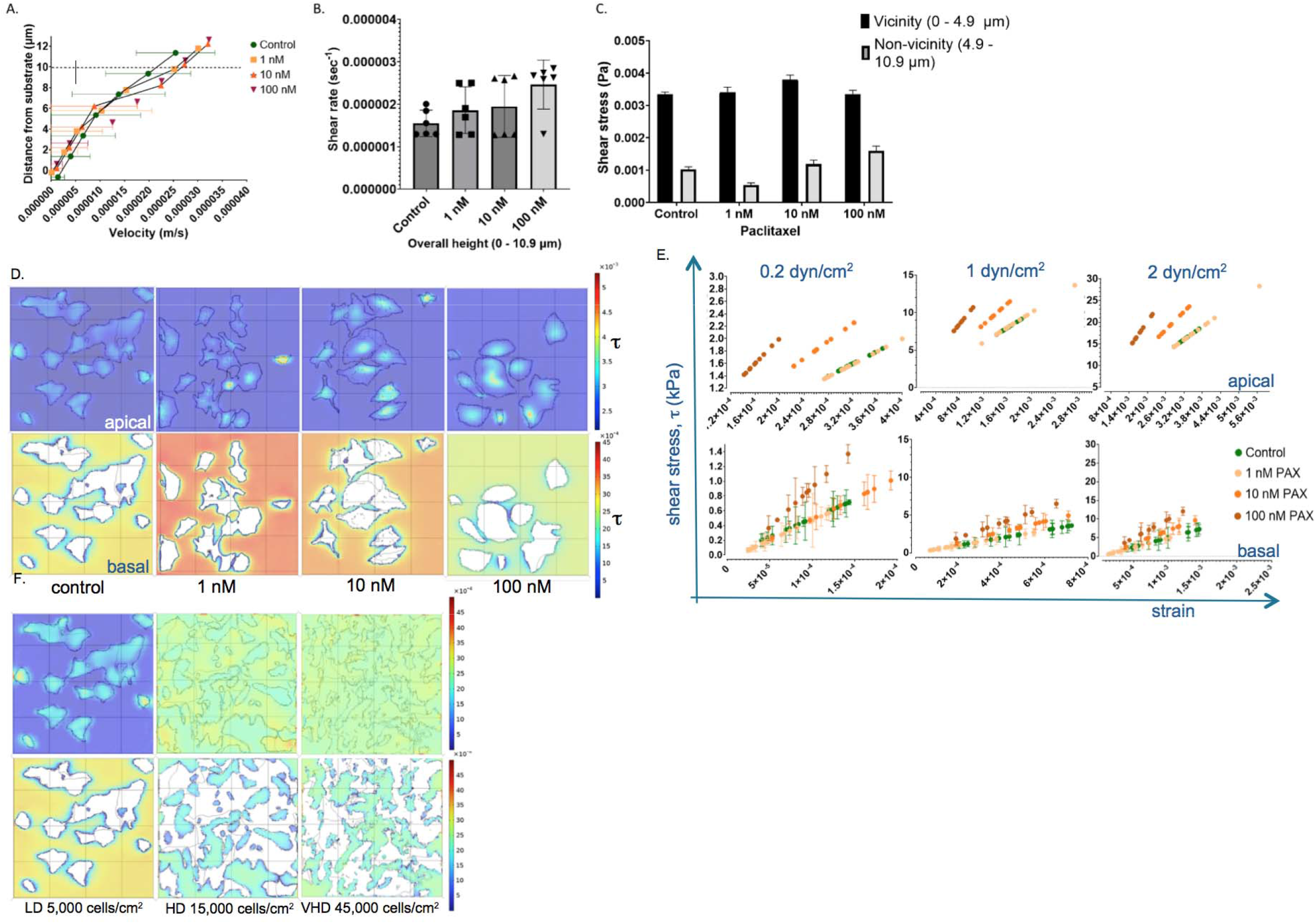
Computational model to elucidate the mechanical milieu of stem cells under flow, expanding upon experimental conditions reported in the current study. Prediction of velocity and shear stress profile on the surface of microtubule-stabilized stem cells (PAX) exposed to flow, which modulates the cytoskeleton while inducing cell shape and volume changes. (A) A shift in velocity profile around stem cells’ most apical region (defined by z=10 μm, compared to most basal region defined by z=0 μm) is predicted as cell stiffening occurs with increasing PAX concentration. (B) Shear rate predictions (linear slope or velocity gradient from (A)) as a function of height, from the base of the chamber to 10.9 µm away from the base of the chamber. (C) Computational predictions of shear stress variation with respect to cell structural changes due to microtubule stabilization (PAX). The wall shear stress calculations show variance between the basal (vicinity) and apical (non-vicinity) region. (D) Computational prediction of shear stress experienced at the apical (top row) and basal (bottom row) region of stem cells across the microtubule-stabilized (PAX) groups (cut through view). (E) The stress-strain curve was plotted using a linear elastic idealization for cell behaviour. With increasing target shear stress magnitude, the shift in the stress-strain slope can be used to map the mechanome prospectively and in the future to guide stem cell differentiation. The shear stress profile predicted for low (F), high (G) and very high (H) density cells in the apical (top row) and basal (bottom row) regions. Flow prediction calculated for a target wall shear stress of 0.2 dyn/cm^2^. Error bars represent the range of minimum and maximum velocity at the defined z-plane (height). Statistical significance is defined by p<0.05.

Control cells exposed to flow showed increased *actb* expression at LD, with flow effects diminishing to nil at higher seeding densities (HD, VHD). Exposure to flow did not affect expression of *tuba1* expression in control cells.

Given the differences attributed to seeding density in both control and microtubule-stabilized cells, we hypothesized that the flow-induced strain on cells, itself influenced by cell seeding density, modulates expression of *actb*.

Time series imaging revealed remodeling of the F-actin and microtubule cytoskeleton over the 60 minutes’ flow exposure, providing quantitative, spatiotemporal measures of stem cell mechanoadaptation. In control cells, F-actin protrusion retracted from the back (away from flow inlet and direction) side (Supplementary Video S5.4, white arrow) and microtubules became more spread as cells optimized their shape and adhered closer to the substrate (Fig. 4B). In microtubule-stabilized cells, long and thick stress fibres remained over time and more actin monomers seemed to be dispersed at later time points. In contrast, the stabilized microtubule bundles shifted position around the nucleus, facilitating cell spreading closer to the substrate (Fig. 4C).

The concentration of F-actin and microtubule per cell was quantified at specific time points, i.e. t_0_, t_30_, and t_60_ minutes’ flow. The labelling of actin and microtubule in live cells using the SPY probes enabled the visualization of endogenous cytoskeleton polymer filaments, as well as cytoskeletal monomers present at the time of incubation. Within the first 30 minutes of flow, F-actin concentration was significantly reduced in control cells; this effect was delayed to 60 minutes in microtubule-stabilized cells (Fig. 4D). Microtubule concentration also decreased gradually with flow, but less significantly in microtubule-stabilized cells (Fig. 4E). In both control and microtubule-stabilized cells F-actin significantly decreased in the apical region over time (Fig. 4F), whereas microtubule decreased less significantly in microtubule- stabilized cells (Fig. 4G). The reduction in both F-actin (Fig. 4H) and microtubule (Fig. 4I) concentration were less significant in the basal region.

Microtubule concentration remained at a high level over 60 minutes in microtubule-stabilized cells. In the front (flow facing) region, F-actin decreased at a higher rate in microtubule- stabilized cells than in control cells (Fig. 4J), whereas in the back (away from flow) region, the decrease in both control and microtubule-stabilized groups was less significant (Fig. 4K). At the front (Fig. 4L) microtubule concentration also maintained a high level with smaller step decreases over time. Similarly in the back (Fig. 4M) the microtubule in both control and the microtubule- stabilized group decreased slowly.

**Spatiotemporal measurement of F-actin and microtubule as a function of flow exposure and microtubule stabilization**. (C) In microtubule-stabilized cells, F-actin exhibited less dynamic behaviour than microtubules which gradually spread as bundles. F-actin and microtubule concentration decreased at different rates, reducing in the first 30 min of flow. F-actin decreased more in microtubule-stabilized cells after 60 minutes’ flow (D) and more significantly than microtubule (E). In the apical region, significant reductio*n* of F-actin occurred gradually, both in control and microtubule-stabilized cells (F), while microtubule concentration remained high in microtubule- stabilized cells at early timepoints (G). In the basal region, both F-actin (H) and microtubule (I) concentration reduced over time but not significantly, and their concentration was maintained at a high level in microtubule-stabilized cells. On cell surfaces facing the flow direction (J) F-actin decreased significantly over time in both control and microtubule-stabilized cells, whereas F-actin on surfaces facing away from the direction of flow did not change significantly (K). The reduction in microtubule in the front (L) and back (M) was gradual but less significant. Error bars represent ± standard error of mean. Significant differences are presented between groups (****p < 0.0001, ***p < 0.001, **p < 0.01, *p < 0.05) and analysed with two-way ANOVA with Tukey’s multiple correlation test.

### Effect of flow and microtubule stabilization on expression of gene markers characteristic of mesenchymal condensation and angiogenesis

Exposure of stem cells to 0.2 dyn/cm^2^ flow for 1h elicited changes in gene expression markers indicative of mesenchymal condensation and angiogenesis. Similar to the mRNA expression of *tuba1* and *actb* reported above, greatest effects of flow were observed in cells seeded at LD. Exposure of LD-seeded control cells to flow resulted in significant upregulation of *msx2* (marker for limb development), with more than 10x upregulation in angiogenesis markers including *sp7*, *vegfa*, *pecam-1*, and *cebpa*. Microtubule-stabilized cells seeded at LD and exposed to flow showed significant eight of the 11 gene markers probed, including of the aforementioned angiogenesis markers plus *col2a1, msx2, pparg* and *acan,* respective markers for skeletogenesis (mesenchymal condensation, later chondrogenesis), limb development, adipogenesis and chondrogenesis (Fig. 5A). Notably, without exposure to flow, microtubule-stabilized cells did not show significant upregulation in two angiogenic markers including *sp7* and *vegfa* or in the adipogenic marker *pparg* or in the chondrogenic marker *acan*.

Seeding stem cells at increased density was associated with less, though still significant upregulation in gene transcription markers. Cells seeded at HD and exposed to flow showed significant downregulation of *col2a1*, *acan*, and *cebpa*. The upregulation of *pecam-1* was reduced in HD seeded cells compared to cells seeded at LD, though still remained significant with microtubule stabilization. At VHD, microtubule-stabilized cells exhibited the highest fold of upregulation in *acan* (3.3-fold), which was abrogated under flow exposure.

### Experimental model - mechanical testing of live stem cells

To better understand mechanical effects, of seeding at increased cell density and exposure to flow, on stem cell architecture as well as protein and gene expression, we created stress – strain plots using our experimental data, akin to a mechanical test of live stem cells as they adapt, and as emergent lineage commitment unfolds. We hypothesized that these differences in normal and shear stress magnitude will be transduced to the nucleus via the cytoskeleton, resulting in differences in lineage specific gene expression.

Shear (Fig. 5B) and normal (Fig. 5C) stress - strain plots were created based on xy- and respective xz-displacements of microbeads attached to stem cells seeded at LD, HD and VHD. In contrast to locally significant differences in strains measured on surfaces of stem cells seeded at increased density and exposed to flow (Fig. 5G-L), mean (global) shear and normal strains showed no significant differences (data not shown).

The increase in both *actb* and *tuba1* mRNA observed with exogenous microtubule stabilization was consistent with the increase of F-actin and microtubule in the polymerized state, quantified from confocal microscopy data (Fig. 5F). Since the concentration and spatial distribution of F-actin and microtubule define cytoskeletal architecture, we then correlated emergent cell architecture to mRNA expression markers of lineage commitment, *e.g.* mesenchymal condensation, adipogenesis, angiogenesis, osteogenesis and chondrogenesis.

### Experimental model – correlation analysis of cytoskeleton and differentiation gene markers

Cytoskeletal architecture, as measured by spatial distribution of F-actin and microtubule concentrations, *i.e.* in the apical *versus* basal and front (flow facing) *versus* back (downward from flow) regions of the cells, showed significant correlation to mRNA measures of emergent lineage commitment. The apical concentration of F-actin correlated significantly and positively with *col2a1*, *vegfa* and *cebpa* expression, whereas basal F-actin correlated significantly and positively with *pecam-1* expression. Similarly, the apical concentration of microtubule correlated significantly and positively with *col2a1, vegfa*, and *cebpa, in addition to acan.* The basal concentration of microtubule showed positive correlation with *col2a1, acan* and *cebpa*, as well as *runx2*, and *pecam-1* (Table 2).

**Table 2:** Multivariate correlation analysis of the F-actin and microtubule cytoskeleton data quantified from confocal imaging (protein level) and the various differentiation gene expression changes of cells seeded at 5000 cells/cm^2^ and exposed to 0.2 dyn/cm^2^ flow for 60 minutes. Cytoskeleton expression levels are based on quantification of F-actin and microtubule live cell images. Green indicates positive correlation and red indicates negative correlation. Grey elements indicate no statistically significant correlations.

|  |  | *runx2* | *col1a1* | *msx2* | *col2a1* | *sox9* | *pparg* | *acan* | *sp7* | *vegfa* | *pecam1* | *cebpa* |
| --- | --- | --- | --- | --- | --- | --- | --- | --- | --- | --- | --- | --- |
| Actin | TOTAL | 0.174 | -0.001 | 0.239 | -0.012 | 0.255 | 0.257 | 0.090 | 0.150 | -0.024 | 0.042 | 0.124 |
|  | apical | 0.127 | 0.199 | -0.092 | 0.326* | 0.056 | -0.017 | 0.282 | 0.168 | 0.516** | 0.279 | 0.357* |
|  | basal | 0.108 | 0.116 | -0.013 | 0.153 | 0.180 | 0.124 | 0.117 | 0.092 | 0.175 | 0.351* | 0.137 |
|  | front | -0.093 | -0.031 | -0.016 | -0.160 | -0.164 | -0.066 | -0.156 | -0.045 | -0.150 | -0.194 | -0.093 |
|  | back | -0.123 | 0.012 | -0.224 | -0.042 | 0.026 | -0.110 | -0.070 | -0.180 | 0.041 | 0.091 | -0.100 |
| Microtubule | TOTAL | 0.069 | 0.085 | -0.042 | 0.159 | 0.135 | 0.032 | 0.137 | 0.066 | 0.283 | 0.225 | 0.178 |
|  | apical | 0.226 | 0.182 | -0.121 | 0.348* | 0.061 | -0.034 | 0.301* | 0.143 | 0.513** | 0.276 | 0.344* |
|  | basal | 0.293* | 0.186 | 0.192 | 0.300* | 0.259 | 0.290 | 0.340* | 0.286 | 0.288 | 0.368* | 0.350* |
|  | front | 0.087 | 0.168 | -0.032 | 0.126 | 0.103 | 0.041 | 0.160 | 0.038 | 0.232 | 0.121 | 0.147 |
|  | back | 0.067 | -0.017 | -0.009 | 0.158 | -0.073 | -0.047 | 0.060 | 0.147 | 0.181 | 0.107 | 0.150 |

\*\*, Correlation is significant at the 0.01 level (2-tailed).
\*, Correlation is significant at the 0.05 level (2-tailed).

Measuring *actb* and *tuba1* at the mRNA level provided a measure of respective gene expression agnostic to cell architecture but in context of changes in gene expression representative of early lineage commitment. When probed at the mRNA level, *actb* expression correlated significantly and positively with that of *col1a1*, *col2a1*, *sox9*, *pparg*, *sp7*, and *vegfa*. (Table 3). In cont*r*ast *tuba1* mRNA expression correlated significantly and positively with *col1a1* and *cebpa*.

**Table 3:**
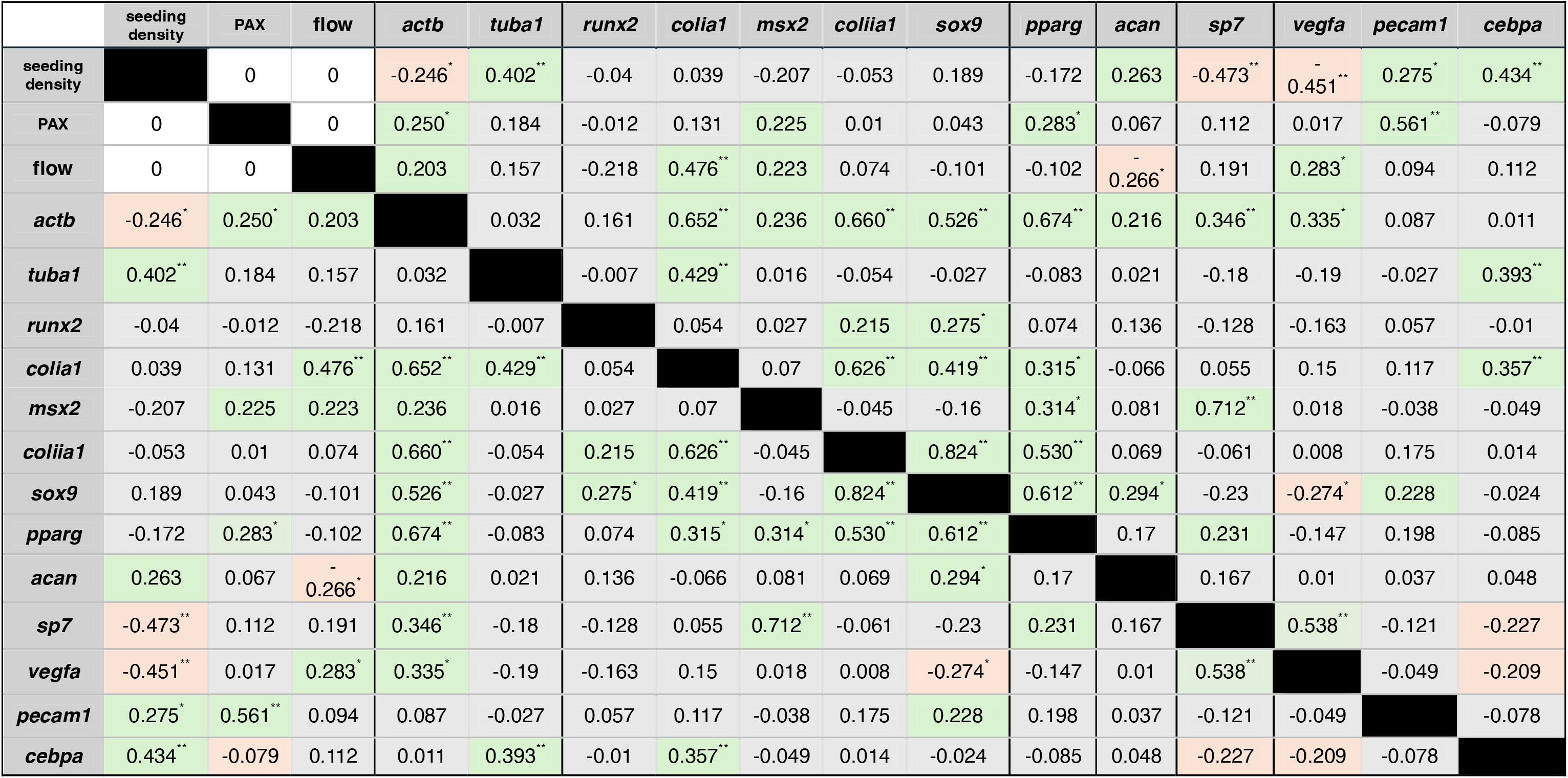
Multivariate correlation analysis for all independent variables, *e.g.* exogenous microtubule stabilization (pax), seeding density, and flow exposure; the actin and microtubule mRNA, actb and tuba1 genes respectively, **and the expression of gene markers for pre-, peri, and post- mesenchymal condensation, osteogenesis, adipogenesis, chondrogenesis, and angiogenesis.** Expression levels are based on the ΔCt values after normalization to Ct values of tbp as the housekeeping gene. Green indicates positive correlation, red indicates negative correlation, and white indicates no correlation. Grey elements indicate no statistically significant correlations. *Correlation signifcant at 0.05 level (2-tailed). **Correlation signifcant at 0.01 level (2-tailed),

Statistical testing of the interactions between each of the genes probed with each independent variable revealed how the environmental (seeding density, exposure to flow) and exogenous (microtubule stabilization) cues independently influenced individual genes. Increased seeding density correlated significantly and positively with *tuba1*, *pecam1*, and *cebfa*, while significantly and negatively with *actb, sp7*, and *vegfa.* Similarly, exogenous microtubule stabilization correlated significantly and positively with *pecam1* but also with *actb* and *pparg*. Interestingly, exposure to flow showed opposite effects to increasing seeding density, with a significant negative correlation to *acan* expression and significant positive correlation to *vegfa* expression, in addition to a significant positive correlation to *col1a1 (*which was not observed to correlate significantly to either seeding density or exogenous microtubule stabilization (Table 3).

To test the influence of the interaction between the independent variables on the expression of *actb* and *tuba1,* between subject effects testing was performed from the three-way ANOVA (Table S3). At a significance level of p<0.05, seeding density significantly influenced *tuba1* expression, whereas exogenous microtubule stabilization (PAX) significantly influenced *actb* expression. No other independent variable interactions significantly influenced *actb* or *tuba1* expression. Seeding density alone significantly influenced the expression of pre- (*runx2, msx2*), and post-mesenchymal condensation markers (*col2a1*, *sox9*). The interaction of all three independent variables, *i.e.* seeding density, exogenous microtubule stabilization (PAX) and flow, influenced *col1a1* (Table S4). Seeding density exerted a significant influence on *vegfa*, *pecam-1* and *cebpa* expression. More importantly, *pecam-1* was highlighted as significantly dependent on the interaction of both seeding density and microtubule stabilization (PAX), or seeding density and flow, as well as interaction of all three variables (Table S5). Interestingly, seeding density significantly influenced, *ppag* and *sp7* expression, respective markers for adipogenesis and osteogenesis, but not *acan*, a marker for chondrogenesis. Furthermore, exogenous microtubule stabilization (exposure to PAX) alone correlated significantly *pparg* expression. Combined with flow, seeding density modulated *acan* and *sp7* expression (Table S6).

### Computational studies to elucidate the mechanical milieu of stem cells under flow

Since our experimental outcome measures revealed strong local effects, of flow interacting with exogenous microtubule stabilization, on stem cell remodeling and emergent lineage commitment, we paired our experimental studies with computational models to elucidate the local mechanical *milieux* of cells. We used a previously published paired computational-experimental approach from our group, since this computational model had been validated experimentally with particle image velocimetry (PIV) experiments^14,17^. Using this paired computational-experimental approach enabled us to, on the one hand, understand local effects of laminar flow influenced by the presence and density of cells themselves, while on the other hand to understand the effects of changes in cell material properties incipient to exogenous microtubule stabilization (PAX). Furthermore, it enabled us to predict virtually the effects of, e.g. increasing flow magnitude and increasing PAX concentrations, on the local mechanical environment of cells, thus expanding on current experimental results.

First, we simulated flow over microtubule-stabilized cells using actual cell geometries from confocal images, and applying flow specifications from the ProFlow chamber as used in the experimental studies (Fig. S3). Experimentally, we had observed that microtubule stabilization results in both gross cell shape and stiffness changes as well as increased cell volume, in a time and concentration-dependent manner^25,26^. In the current experimental study, we observed ConA bead-tagged cells, to show less bead displacement, *i.e.* cells behaved more rigidly, deforming less under flow than control cells. Computational simulations, in microtubule-stabilized cell models incorporating experimental observations of larger cell volume and flatter structure, predicted small increases in flow velocity (Fig. 6A) and shear rate (Fig. 6B) toward the apical surface of the cell. Furthermore, significant differences in shear stress were predicted between the apical (non-vicinity) and basal (vicinity) regions of control and PAX-treated cells (Fig. 6C).

We then used coordinates in both the apical (apical most point) as well as the basal (three selected coordinates) regions of the cells as reference points (Fig. 6D) to plot the stress-strain curve of the microtubule-stabilized cells using the experimentally measured (AFM) Young’s modulus; stress – strain curves created in this way consistently demonstrated a shift in the slope and intercept, indicative of the stiffening effect of microtubule stabilization (PAX). Interestingly, computational models predicted non-intuitive effects of interactions between flow and cell stiffening due to microtubule stabilization; namely, cell stiffening was predicted to be flow-rate dependent, albeit with **opposite** effects on the slope for increasing flow velocities, i.e. stiffening (higher slope) in apical regions with increasing flow velocity and softening (lower slope) in basal regions with increasing flow velocity (Fig. 6E). Similar to predictions with untreated controls, the largest and most significant predicted differences in shear stresses were between the apical and basal regions of cells seeded at low-density, where the apical surfaces are not buffered by the tight packing of neighbouring cells (Fig. 6F, G, H).

With increasing seeding density, the velocity and shear rate profile were predicted to vary significantly across the height of the cells (Fig. S5A). In areas towards the apical region of the cell (non-vicinity, 4.9–10.9 μm above the substrate), significant differences in shear stress were predicted between the low density (LD) group (Fig. S5B, C). Meanwhile, no differences were predicted between the apical and basal region for the higher density groups, i.e. high density (HD) and very high density (VHD) (Fig. S5D, E). With VHD - 45,000 cells/cm^2^ density mimicking physiological conditions including mesenchymal condensation or formation of multicellular constructs, the volumetric view of shear stress distribution in the sub-model reveals how the higher seeding density influences the mechanical milieu across the thickness of the cell, but negligible when across the thickness of the chamber (Fig. S5F, G,H).

## Discussion

We elucidated the effects of exogenous microtubule stabilization (PAX) on stem cells’ capacity to sense and adapt to changes in their local mechanical environment. Using previously developed experimental and computational tools for respective delivery and prediction of shape and volume changing stresses^11,14,17^, we studied the interplay between the living, evolving cells and their mechanical environment, and tested specific hypotheses as detailed below. The significant changes in MSCs’ material properties (stiffened) and volume (increased) induced by exogenous microtubule stabilization resulted in significantly different normal and shear stress loading histories compared to control cells exposed to identical bulk laminar flow (0.2 dyn/cm^2^). Using a paired computational model, we further predicted a range of mechanoadaptation responses of microtubule-stabilized cells to scaled up flow magnitudes (1 and 2 dyn/cm^2^). In experimental studies, these spatiotemporal mechanical cues transduced to the nucleus via the cytoskeleton, triggering changes in gene expression indicative of emergent lineage commitment. Exogenous modulation of the MSC cytoskeleton changed not only the cells’ capacity to sense and adapt to changes in their mechanical *milieux*, but also resulted in significantly different changes in gene expression indicative of early lineage commitment, compared to those observed in naive (no microtubule stabilization) control cells. Hence, MSCs adapt to as well as modulate their own mechanical environment via cytoskeletal remodeling and lineage commitment - microtubule stabilization changes not only MSCs’ mechanoadaptive machinery, their capacity to adapt, and their lineage commitment, but also their mechanical environment. Taken as a whole, these studies corroborate our working hypothesis that MSCs and their mechanoadaptive machinery serve as sensors and actuators, intrinsically linked to their lineage potential via mechanoadaptive feedback loops which are sensitive to exogenous modulation via biochemical and biophysical means^2^.

Previous studies have demonstrated that stem cells exhibit exquisite sensitivity to changes in their mechanical *milieux*. Compared to terminally differentiated cells, stem cells respond to volume- and shape changing stresses of 1000x smaller magnitude, eliciting changes in their cytoskeleton and up-/down-regulating gene expression indicative of emergent lineage commitment^1,2,8,9,12,13^. In terminally differentiated cells, adaptation to physiological flow plays a role in how cells maintain tissue homeostasis, *e.g.*, maintenance of epithelial integrity and barrier function^28^, and vascular endothelial hemodynamic function^29^. This adaptation involves changes in membrane fluidity, cell adhesion, cytoskeleton, and collective cell migration^30^. Cells with perturbations in any protein component of cellular function, *e.g.*, the glycocalyx^29^, lose their capacity to adapt to flow. Similarly, as observed in the current study, exogenous perturbation of the microtubule influences the cells‘ homeostatic capacity. Compared to control cells, microtubule-stabilized cells showed smaller magnitude microbead displacements in the xy-direction under laminar flow, likely due to the stiffening effect of PAX. In addition, laminar flow is known to elevate tension in the actin cytoskeleton^31^, which could intensify the effect of PAX in cytoskeletal stabilization. As a whole, these observations point to new avenues to guide stem cell mechanoadaptation and targeted lineage commitment.

Our approach to map stress-strain and incipient mechanoadaptation enabled mechanical testing and quantification of **spatiotemporal adaptation of live stem cells**, a distinct advantage compared to single time point and/or single cell testing of mechanical properties using AFM or deformability cytometry.^32^ In the current study, bead displacements were used as markers of flow-induced displacements at cell-fluid interfaces. This idealization does not account for the traction generated by the cellular actin-myosin system^26^ which is itself directly impacted by stabilization of microtubules. Using our methods, cells were observed to adapt immediately after exposure to flow, reducing their height, and effectively reducing drag at cell-flow interfaces. Seeding microtubule-stabilized cells at increased density mitigated both flow-induced strains as well as mechanoadaptation of cells via cytoskeletal protein expression and remodeling, and incipient changes in gene expression indicative of emergent lineage commitment. It has been reported previously that microtubule-stabilized cells seeded at increasing seeding density exhibit a higher increase in bulk modulus compared to control cells^25,26^. In the current study, microtubule-stabilized cells seeded at LD exhibited less dynamic displacement in the z-direction than control cells which gradually decreased their height over time (less prominent, less flow disruption). At HD and VHD, both control and microtubule-stabilized cells exhibit dynamic displacement in the z-direction, suggesting that as multicellular construct they collectively adapted to match the flow conditions regardless of cytoskeletal perturbations. Previous reports indicate an important modulatory role for mechanosensitive ion channels in cytoskeletal remodeling where Ca^2+^ ion channel inhibitors, Gd3+ and GsMTx4, eliminated flow-induced actin reorganization^33^.

Hence, dynamic cytoskeletal remodeling forms one cellular mechanoadaptation response, where deformation drives stiffening or softening of cells (Young’s modulus). Previously reported AFM measurements demonstrate changes in cell stiffness (Young’s modulus) with microtubule stabilization (PAX), seeding at increased density, and culture on compliant substrates, all indicative of mechanoadaptation^25,26^. Similarly, cell deformations, *i.e.* the shear and normal strains measured as cell surface displacements under the exposure to flow or controlled mechanical loads, can be used to predict the Young’s modulus of cells *in situ,* in tissue templates or multicellular constructs, and increasingly in tissues themselves^34^. A current limitation of the approach is the idealization of cells as linear elastic; new mechanics approaches such as the virtual power theory will increasingly enable a more realistic elucidation of cell and tissue mechanics^2^. These new approaches will also expand our understanding as cells proliferate and evolve into multicellular constructs, enabling bottom-up studies, from molecular levels upwards of spatiotemporal cytoskeletal remodeling and its functional implications at longer length and time scales.

Actin and microtubule cytoskeleton remodeled in response to flow exposure, and in interaction with microtubule stabilization and cell seeding density. Interestingly, immediately after exposure to flow, microtubule-stabilized cells exhibited higher and more significant increases in F-actin and microtubule protein expression than control cells; these changes decreased with time, indicative of adaptation. Furthermore, spatiotemporal adaptation of cytoskeletal F-actin and microtubules was observed as a flow-direction dependent change in architecture, *i.e.* with most significant changes in the flow facing front *versus* the slip stream back half of the cell. Taken together, microtubule stabilization stiffens stem cells, which reduces strains induced by flow (bead displacements); in contrast, microtubule stabilization increases the expression of cytoskeletal proteins and changes their spatiotemporal distribution in a flow-direction dependent manner. In essence, although less deformed by flow, microtubule-stabilized cells appear to be more mechanoadaptive, though detailed temporal scaling of protein expression and architectural remodeling are needed to elucidate specific effects elicited by microtubule stabilization versus flow-induced remodeling.

Elucidation of temporal differences in cytoskeleton gene (expression and post-translational modifications) *versus* protein level (expression, de-/polymerization) mechanoadaptation will be key to deciphering and harnessing underpinning mechanisms in context of both embryonic development and regenerative medicine. For example, at the gene level, microtubule stabilization is associated with higher expression of *tuba1* compared to *actb* (gene level). In contrast, at the protein level, microtubule stabilization results in increased polymerization of both microtubule and, to a higher degree, F-actin (protein level)^25,26^. These temporal differences can be further parsed into spatial differences, *e.g.* the protein level increase in F-actin is more prominent in the basal region of the cell, while the increase in microtubule is more global across both apical and basal regions. These differences may reflect different time scales in gene and protein level changes, *e.g.* the duration of microtubule exposure to PAX versus the longer time scale required for cells to maintain adhesion to the glass substrate over the course of the experiment. Furthermore, at the molecular level, stabilization of microtubules requires acetylation of α-tubulin, an important post-translational modification (PTM) which limits the flexibility of the inside of microtubule loop^35^. A functional consequence of the loss of α-tubulin acetylation in development represents a hallmark for epithelial-to-mesenchymal transition (EMT), where cytoskeletal remodelling occurs dramatically as F-actin accumulates at the leading edge, promoting cell polarity and migratory phenotype^36^. In context of our current experiments, exposure to flow with microtubule stabilization could emulate similar processes, compensating for the microtubule and promoting the increase in *actb* instead of *tuba1*. Hence, change in the mechanical *milieux* of cells can immediately modulate cytoskeletal adaptation and thereby control the rate of actin and microtubule de-/polymerization to a greater extent than modulation via chemical agents.

In the current study we observed that differences in normal and shear stress magnitude that were transduced to the nucleus via the cytoskeleton, resulting in differences in lineage specific gene expression. This observation corroborates previous published studies using a range of MSC types (human, rat, sheep, and canine derived), and a range of flow regimes, *e.g.* 1 dyn/cm^2^ to 15 – 25 dyne/cm^2^ for 12 hours to 14 days, has been shown to enhance the upregulation of endothelial cells (ECs) markers including CD31, von Willebrand Factor, and vascular endothelial-cadherin (VE-cadherin)^37^. Shear-stress exposure has been reported to upregulate platelet/endothelial cell adhesion molecule 1 (PECAM-1) in mouse embryonic stem cells (ESCs) and human adipose tissue-derived stem cells (ASCs).^37^ In the current study, we found that microtubule stabilization alone (without flow) significantly upregulated *pecam-1.* This significant increase in *pecam-1* was maintained with exposure to flow. At higher seeding densities we observed upregulation of *vegfa* and no upregulation in *pecam-1.* The differences in angiogenic marker expression may reflect the different biological timelines of angiogenic marker expression in healthy and pathologic development^38,39^, *e.g. pecam-1* mostly plays a role in angiogenesis during early development and tumour metastasis in disease, while *vegfa* plays a role in maintenance of mature blood vessels^40^.

Expanding upon previous studies delivering minute mechanical cues relevant to *in utero* development^13,14^, we observed similar anisotropic changes in actin and microtubule concentrations and spatial distributions. At the gene level, actin was significantly affected by microtubule stabilization and flow exposure, whereas microtubule was affected by seeding density. Changes in actin concentration at protein level, especially in the apical region, consistently correlated with the *coliia1* gene, and actin at gene level (*actb*) correlated positively with more gene markers for mesenchymal condensation, osteogenesis, chondrogenesis, and angiogenesis. The observed increase in microtubule at apical and basal regions of the cells correlated positively with markers of mesenchymal condensation and chondrogenesis. In contrast, at the gene level *tuba1* correlated with fewer genes, *e.g. coliia1* and *cebpa*. This anisotropy reflects the distinct mechanical function of actin and microtubule at various timepoints of development as differentiating cells balance forces in condensing and proliferating tissues (higher density) and in response to dynamic normal and shear stresses from *e.g.* deformation and flow.

The strong effects of seeding density and time in culture^1,7–10^, in particular on epigenetic mechanisms and differentiation of stem cells (hiPSCs, towards ectoderm, mesoderm, or endoderm lineage)^38^ are well documented. As cells transition from exponential to stationary phase at 72h to 120h culture time, stem cells exhibit increasing ratio of H3K4me3/H3K27me3, the respective activating and repressing histone modification, that corresponds to high expression of pluripotency markers, *oct4*, *nanog*, and *sox2*^41^. Culture of hiPSCs into high density pellets consistently promoted chondrogenesis with strong collagen II staining^42^. Transfer of those pellets to rotary culture to introduce shear stress further enhanced cartilage tissue containing larger chondrocytes, indicative of hypertrophic chondrogenesis^42^. In line with transcriptional changes in early development, single cell RNA sequencing on human epiphyseal cells isolated from 12-32 weeks post-conception enabled identification of temporal switch in ossification and trajectories of various populations commitment with distinct collagen expression.^43^ Similarly, across the LD, HD, and VHD cells exposed to flow in this study, genes that were significantly upregulated in LD, such as *col2a1* and *acan*, were downregulated in HD. At VHD, these cartilage-associated markers were upregulated again, suggesting that emulating the stages of multicellular construct relevant to the timeline of development could reveal a similar temporal genetic change. Combined mechanomics-transcriptomics could be a useful strategy towards understanding the temporal genetic changes during development. Further investigations of stem cell adaptation in longer-term culture and flow conditions as well as utilizing *e.g.* embryoid bodies, to model specific tissues could provide further information on cell and multicellular construct mechano-responses across the relevant developmental timeline.

Our paired computational – experimental model enabled us to expand upon our experimental results, *e.g.* to predict effects of decreased microtubule stabilization (lower PAX concentration), and higher flow magnitudes, on the mechanical behavior of cells (stress-strain plots). Computational flow simulations revealed that the presence cells and their modulated structures upon PAX treatment contribute to the varying velocity and shear rate across the height of the cells. Microtubule stabilization with 1 nm PAX resulted in no significant changes to mechanical behavior of cells for flows at 0.2, 1 and 2 dyn/cm^2^. Microtubule stabilization with 10 nm PAX resulted in stress-strain plots intermediate to those of the 1 and 100 nm PAX microtubule-stabilized cells. Shear rate, the rate of change in velocity across the apical-basal height of cells, increased around the cells with increasing PAX concentration, which is contributed by the changing cellular shape related to flattening and remodelling of topography of cell surface. By tuning exogenous cytoskeletal de-/polymerization, together with delivery of conducive mechanical cues, future smart materials and devices may provide new approaches to guide spatiotemporal targeted tissue genesis in engineered tissue templates. The stiffening effect of PAX contributes to variations in local stresses as well as stress-strain behaviour of cells, and thus adds complexity to elucidation of the emergent mechanical milieu. While our current simulation was performed with the assumption that the stress components related to strain by the elastic modulus, it is important to note that future studies should consider the time dependence or the deformation that cells would exhibit as a function of the loading rates^44^. The current predictive study depicts an instantaneous stress acting on the cell and not how fast or for how long the flow is introduced. In the future it will be important to model potential non-linear elastic properties such as stiffening and softening, in addition to potential inelastic *i.e.* viscoelastic, viscoplastic, and poroelastic properties. Furthermore, the adaptive behaviour of cells makes the mapping of a fully realistic mechanome particularly challenging, *e.g.* including capturing the evolution of stem cell mechanoresponses emulating developmental stages, and postnatal healing.

Collective multidisciplinary work integrating mechanical and biophysical cues conducive to guiding specific lineage commitment^45^, while retrospectively incorporating insights from mechanoadaptation studies in terminally differentiated cells, with new *in vitro*, *in silico*, and *in situ* platforms, is key to mapping and leveraging the mechanome. Ultimately, we aim to create functional tissue templates that can perform in physiologically relevant contexts, enabling the controlled delivery of mechanical, biophysical and biochemical cues,^46^ in both health and disease.

## Conclusion

The adaptive response of stem cells to their dynamic mechanical environment represents a key driver of their structure and function relationships as they proliferate during the processes of tissue development and postnatal healing. Mapping of the mechanical milieu and corresponding subcellular (cytoskeletal, gene expression) and cellular responses, *i.e.* the mechanome, presents a valuable, benign (compared to *e.g.* gene editing) platform to guide stem cell lineage commitment, guiding targeted tissue neogenesis using mechanical cues for regenerative medicine. Yet successful mapping of the mechanome will require both top-down and bottom-up strategies enabling the study and elucidation of multiscale cellular adaptation in context of complex physiological processes of development and postnatal healing. To address this challenge, novel tools have been developed and validated to enable controlled delivery of mechanical cues relevant to those in development and to visualize and measure stem cell adaptation as it occurs^5^. In this study, we present the application of paired live cell experimental and predictive computational approaches to probe effects of controlled volume- and shape-changing stress (*e.g.*, compression and shear respectively) on MSC structure and function and emergent lineage commitment^11,14^. By stabilizing the microtubule cytoskeleton with paclitaxel (PAX)^47,48^, we could probe stem cell mechanoadaptation in naive control cells and cells with compromised mechanoadaptation machinery^25,26^. The study opens new possibilities for tuning mechanical *milieux* and their cellular inhabitants to guide mechanoadaptation, *i.e.* cell structure and emergent function, to perform in targeted contexts.

This study highlights the novel methods for mapping the mechanome previously reported in Song et al.2012^14^ and Chang and Knothe Tate 2011^13^, through controlled delivery of volume- and shape-changing stresses by increasing seeding density (compression) and introducing laminar flow to induce shear stress respectively. Experimental flow exposure to cells seeded and various densities, while visualizing and measuring cell structural changes over time, presents a powerful tool to study stem cells time-dependent adaptation capacity that govern their mechanical properties. Spatiotemporal analysis of stem cells adaptation to stresses relevant to those in development, while modulating the cytoskeleton using chemical agents, enables the study of stem cells immediate mechanoresponses through regulation of shape, volume, actin and microtubule remodeling and spatial distribution that can be correlated with the expression of gene markers for tissue specific differentiation. The anisotropy of actin and microtubule was also confirmed with their distinct rate of change in concentration during adaptation to high seeding density and flow exposure, and when perturbed with PAX. These responses such as cytoskeletal remodelling, focal adhesion enrichment, and gradient of other mechanosensitive proteins, as well as the transcripts of their genome^49^, could provide an important references library and a powerful -omics analysis tool e.g. the mechanomics, for studying system biology and targeting stem cells differentiation using mechanical cues. F-actin and microtubule also exhibit contrasting trend in concentration changes between polymerized state and at mRNA level, demonstrating the extent of effects of chemical or mechanical cues on the cytoskeleton. Overall, exposure to flow and increasing seeding density, together with the use of PAX to inhibit the function of microtubule presents a novel tool for controlled delivery of cues towards further understanding the mechanism of stem cells and cytoskeletal adaptation across scales of tissue development and regeneration, and the role of interplaying mechanical and biochemical cues in guiding stem cells differentiation. Future work on expanding the library of stem cells mechanical properties and how they evolve over time using these mechanomics engineering tools, along with their differentiation responses at various stages of development, would enable the refinement of the mechanome map and future studies aiming to target stem cells fate or tissue neogenesis for tissue engineering and regenerative medicine.

## References

1. M. L. Knothe Tate, P. W. Gunning, and V. Sansalone, “Emergence of form from function—Mechanical engineering approaches to probe the role of stem cell mechanoadaptation in sealing cell fate,” Bioarchitecture, vol. 6, no. 5, pp. 85– 103, Sep. 2016.

2. M. L. Knothe Tate, T. D. Falls, S. H. McBride, R. Atit, and U. R. Knothe, “Mechanical modulation of osteochondroprogenitor cell fate,” International Journal of Biochemistry and Cell Biology, vol. 40, no. 12. pp. 2720–2738, 2008.

3. T. Mammoto and D. E. Ingber, “Mechanical control of tissue and organ development,” Development. 2010.

4. P. Patwari and R. T. Lee, “Mechanical Control of Tissue Morphogenesis,” Circ Res, vol. 103, no. 3, pp. 234–243, Aug. 2008.

5. V. D. L. V. D. L. Putra et al., “Mechanomics Approaches to Understand Cell Behavior in Context of Tissue Neogenesis, During Prenatal Development and Postnatal Healing,” Frontiers in Cell and Developmental Biology, vol. 7. 2020.

6. E. J. Anderson, T. D. Falls, A. M. Sorkin, and M. L. Knothe Tate, “The imperative for controlled mechanical stresses in unraveling cellular mechanisms of mechanotransduction.,” Biomed Eng Online, vol. 5, p. 27, May 2006.

7. M. L. Knothe Tate, T. D. Falls, S. H. McBride, R. Atit, and U. R. Knothe, “Mechanical modulation of osteochondroprogenitor cell fate,” International Journal of Biochemistry and Cell Biology. 2008.

8. S. H. McBride and M. L. Knothe Tate, “Modulation of stem cell shape and fate A: The role of density and seeding protocol on nucleus shape and gene expression,” Tissue Eng Part A, vol. 14, no. 9, pp. 1561–1572, 2008.

9. S. H. McBride, T. Falls, and M. L. Knothe Tate, “Modulation of stem cell shape and fate B: Mechanical modulation of cell shape and gene expression,” Tissue Eng Part A, vol. 14, no. 9, pp. 1573–1580, 2008.

10. V. D. L. Putra, K. A. Kilian, and M. L. Knothe Tate, “Biomechanical, biophysical and biochemical modulators of cytoskeletal remodelling and emergent stem cell lineage commitment,” Commun Biol, vol. 6, no. 1, p. 75, 2023.

11. E. J. Anderson and M. L. Knothe Tate, “Open access to novel dual flow chamber technology for in vitro cell mechanotransduction, toxicity and pharamacokinetic studies,” Biomed Eng Online, vol. 6, no. 1, p. 46, 2007.

12. M. J. Song, D. Dean, and M. L. Knothe Tate, “Mechanical modulation of nascent stem cell lineage commitment in tissue engineering scaffolds,” Biomaterials, 2013.

13. Hana. Chang and M. L. Knothe Tate, “Structure - Function relationships in the stem cell’s mechanical world B: emergent anisotropy of the cytoskeleton correlates to volume and shape changing stress exposure,” MCB Molecular and Cellular Biomechanics, vol. 8, no. 4, pp. 297–318, 2011.

14. M. J. Song, S. M. Brady-Kalnay, S. H. McBride, P. Phillips-Mason, D. Dean, and M. L. K. Tate, “Mapping the mechanome of live stem cells using a novel method to measure local strain fields in situ at the fluid-cell interface,” PLoS One, 2012.

15. J. A. Zimmermann and M. L. Knothe Tate, “Structure - Function relationships in the stem cell’s mechanical world A: Seeding protocols as a means to control shape and fate of live stem cells,” MCB Molecular and Cellular Biomechanics, vol. 8, no. 4, pp. 275–296, 2011.

16. S. F. Evans, D. Docheva, A. Bernecker, C. Colnot, R. P. Richter, and M. L. Knothe Tate, “Solid-supported lipid bilayers to drive stem cell fate and tissue architecture using periosteum derived progenitor cells,” Biomaterials, vol. 34, no. 8, pp. 1878–1887, 2013.

17. M. J. Song, D. Dean, and M. L. Knothe Tate, “In situ spatiotemporal mapping of flow fields around seeded stem cells at the subcellular length scale,” PLoS One, 2010.

18. H.-W. Lee, J. H. Shin, and M. Simons, “Flow goes forward and cells step backward: endothelial migration,” Exp Mol Med, vol. 54, no. 6, pp. 711–719, 2022.

19. C. P. Ng and M. A. Swartz, “Fibroblast alignment under interstitial fluid flow using a novel 3-D tissue culture model,” American Journal of Physiology-Heart and Circulatory Physiology, vol. 284, no. 5, pp. H1771–H1777, May 2003.

20. L. Dan, C.-K. Chua, and K.-F. Leong, “Fibroblast response to interstitial flow: A state-of-the-art review,” Biotechnol Bioeng, vol. 107, no. 1, pp. 1–10, Sep. 2010.

21. E. A. Osborn, A. Rabodzey, C. F. J. Dewey, and J. H. Hartwig, “Endothelial actin cytoskeleton remodeling during mechanostimulation with fluid shear stress.,” Am J Physiol Cell Physiol, vol. 290, no. 2, pp. C444–52, Feb. 2006.

22. R. P. Wolfe and T. Ahsan, “Shear stress during early embryonic stem cell differentiation promotes hematopoietic and endothelial phenotypes.,” Biotechnol Bioeng, vol. 110, no. 4, pp. 1231–1242, Apr. 2013.

23. E. J. Anderson and M. L. Knothe Tate, “Design of Tissue Engineering Scaffolds as Delivery Devices for Mechanical and Mechanically Modulated Signals,” Tissue Eng, vol. 13, no. 10, pp. 2525–2538, Sep. 2007.

24. K. A. Kilian, B. Bugarija, B. T. Lahn, and M. Mrksich, “Geometric cues for directing the differentiation of mesenchymal stem cells.,” Proc Natl Acad Sci U S A, vol. 107, no. 11, pp. 4872–4877, Mar. 2010.

25. V. D. L. Putra, K. A. Kilian, and M. L. K. Tate, “Stem cell mechanoadaptation - Part A - Effect of microtubule stabilization and volume changing stresses on cytoskeletal remodeling,” bioRxiv, p. 2024.07.28.605421, Jan. 2024.

26. V. D. L. Putra, K. A. Kilian, and M. L. Knothe Tate, “Stem cell mechanoadaptation Part B - Microtubule stabilization and substrate compliance effects on cytoskeletal remodeling,” bioRxiv, p. 2024.07.28.605537, Jan. 2024.

27. S. Dokos, “Fluid Mechanics,” in Modelling Organs, Tissues, Cells and Devices: Using MATLAB and COMSOL Multiphysics, S. Dokos, Ed. Berlin, Heidelberg: Springer Berlin Heidelberg, 2017, pp. 305–341.

28. T. P. J. Wyatt et al., “Emergence of homeostatic epithelial packing and stress dissipation through divisions oriented along the long cell axis,” Proceedings of the National Academy of Sciences, vol. 112, no. 18, pp. 5726–5731, May 2015.

29. D. A. Chistiakov, A. N. Orekhov, and Y. V Bobryshev, “Effects of shear stress on endothelial cells: go with the flow,” Acta Physiologica, vol. 219, no. 2, pp. 382–408, Feb. 2017.

30. J. A. Espina, M. H. Cordeiro, M. Milivojevic, I. Pajić-Lijaković, and E. H. Barriga, “Response of cells and tissues to shear stress,” J Cell Sci, vol. 136, no. 18, p. jcs260985, Sep. 2023.

31. E. Tzima, “Role of Small GTPases in Endothelial Cytoskeletal Dynamics and the Shear Stress Response,” Circ Res, vol. 98, no. 2, pp. 176–185, Feb. 2006.

32. Y. Hao, S. Cheng, Y. Tanaka, Y. Hosokawa, Y. Yalikun, and M. Li, “Mechanical properties of single cells: Measurement methods and applications,” Biotechnol Adv, vol. 45, p. 107648, 2020.

33. D. Verma, N. Ye, F. Meng, F. Sachs, J. Rahimzadeh, and S. Z. Hua, “Interplay between Cytoskeletal Stresses and Cell Adaptation under Chronic Flow,” PLoS One, vol. 7, no. 9, p. e44167, Sep. 2012.

34. S. H. McBride, S. Dolejs, S. Brianza, U. Knothe, and M. L. Knothe Tate, “Net Change in Periosteal Strain During Stance Shift Loading After Surgery Correlates to Rapid De Novo Bone Generation in Critically Sized Defects,” Ann Biomed Eng, vol. 39, no. 5, pp. 1570–1581, 2011.

35. L. Eshun-Wilson et al., “Effects of α-tubulin acetylation on microtubule structure and stability,” Proceedings of the National Academy of Sciences, vol. 116, no. 21, pp. 10366–10371, May 2019.

36. S. Gu et al., “Loss of α-Tubulin Acetylation Is Associated with TGF-β-induced Epithelial-Mesenchymal Transition *,” Journal of Biological Chemistry, vol. 291, no. 10, pp. 5396–5405, Mar. 2016.

37. Y. Huang, J.-Y. Qian, H. Cheng, and X.-M. Li, “Effects of shear stress on differentiation of stem cells into endothelial cells.,” World J Stem Cells, vol. 13, no. 7, pp. 894–913, Jul. 2021.

38. N. O. Abutaleb and G. A. Truskey, “Differentiation and characterization of human iPSC-derived vascular endothelial cells under physiological shear stress,” STAR Protoc, vol. 2, no. 2, p. 100394, 2021.

39. J. Kroon, N. Heemskerk, M. J. T. Kalsbeek, V. de Waard, J. van Rijssel, and J. D. van Buul, “Flow-induced endothelial cell alignment requires the RhoGEF Trio as a scaffold protein to polarize active Rac1 distribution.,” Mol Biol Cell, vol. 28, no. 13, pp. 1745–1753, Jul. 2017.

40. N. G. dela Paz, T. E. Walshe, L. L. Leach, M. Saint-Geniez, and P. A. D’Amore, “Role of shear-stress-induced VEGF expression in endothelial cell survival.,” J Cell Sci, vol. 125, no. Pt 4, pp. 831–843, Feb. 2012.

41. M.-H. Kim, N. Thanuthanakhun, S. Fujimoto, and M. Kino-oka, “Effect of initial seeding density on cell behavior-driven epigenetic memory and preferential lineage differentiation of human iPSCs,” Stem Cell Res, vol. 56, p. 102534, 2021.

42. S. R. Lamandé et al., “Modeling human skeletal development using human pluripotent stem cells,” Proceedings of the National Academy of Sciences, vol. 120, no. 19, p. e2211510120, May 2023.

43. Z. Deng et al., “Temporal transcriptome features identify early skeletal commitment during human epiphysis development at single-cell resolution,” iScience, vol. 26, no. 8, p. 107200, 2023.

44. W. Jung, J. Li, O. Chaudhuri, and T. Kim, “Nonlinear Elastic and Inelastic Properties of Cells,” J Biomech Eng, vol. 142, no. 10, Aug. 2020.

45. M. L. Knothe Tate, I. Jalilian, M. Jae Song, and S. McBride, “Mapping the Mechanome: New Experimental and Computational Approaches to Elucidate Stem Cell Mechanoadaptation and Lineage Commitment,” Biophys J, vol. 110, no. 3, p. 24a, Feb. 2016.

46. H. Wang et al., “Mechanomics Biomarker for Cancer Cells Unidentifiable through Morphology and Elastic Modulus,” Nano Lett, 2021.

47. M. Kiwanuka et al., “Effect of paclitaxel treatment on cellular mechanics and morphology of human oesophageal squamous cell carcinoma in 2D and 3D environments,” bioRxiv, p. 2022.03.06.483167, Jan. 2022.

48. B. H. Long and C. R. Fairchild, “Paclitaxel inhibits progression of mitotic cells to G1 phase by interference with spindle formation without affecting other microtubule functions during anaphase and telephase.,” Cancer Res, vol. 54, no. 16, pp. 4355–4361, Aug. 1994.

49. J. Wang, D. Lü, D. Mao, and M. Long, “Mechanomics: An emerging field between biology and biomechanics,” Protein and Cell. 2014.

