## Supplementary Information Text and Figures for "Tuning mechanical *milieux* of tissue templates and their cellular inhabitants to guide mechanoadaptation"

Vina D. L. Putra^1^, Vittorio Sansalone^2^, Kristopher A. Kilian^1*^, Melissa L. Knothe Tate^3*^

^1^School of Chemistry and School of Materials Science & Engineering, University of New South Wales, Sydney, NSW, Australia ^2^Laboratoire Modélisation et Simulation Multi Echelle, Equipe Biomécanique, Faculté des Sciences et Technologie, Université Paris-Est Créteil Val de Marne, France

^3^Blue Mountains World Interdisciplinary Innovation Institute (bmwi³), Blue Mountains, NSW, Australia

ORCID for MLKT: 0000-0002-3552-2525

**Classification:** Biological Systems Engineering, Computational Simulations, Cell Biology, Biophysics

**Keywords:** Mechanoadaptation, Remodeling, Stiffness, **C**ytoskeleton, Development, Computational Fluid Dynamics Modeling

**Supplementary Information**

*EXPERIMENTAL FLOW*

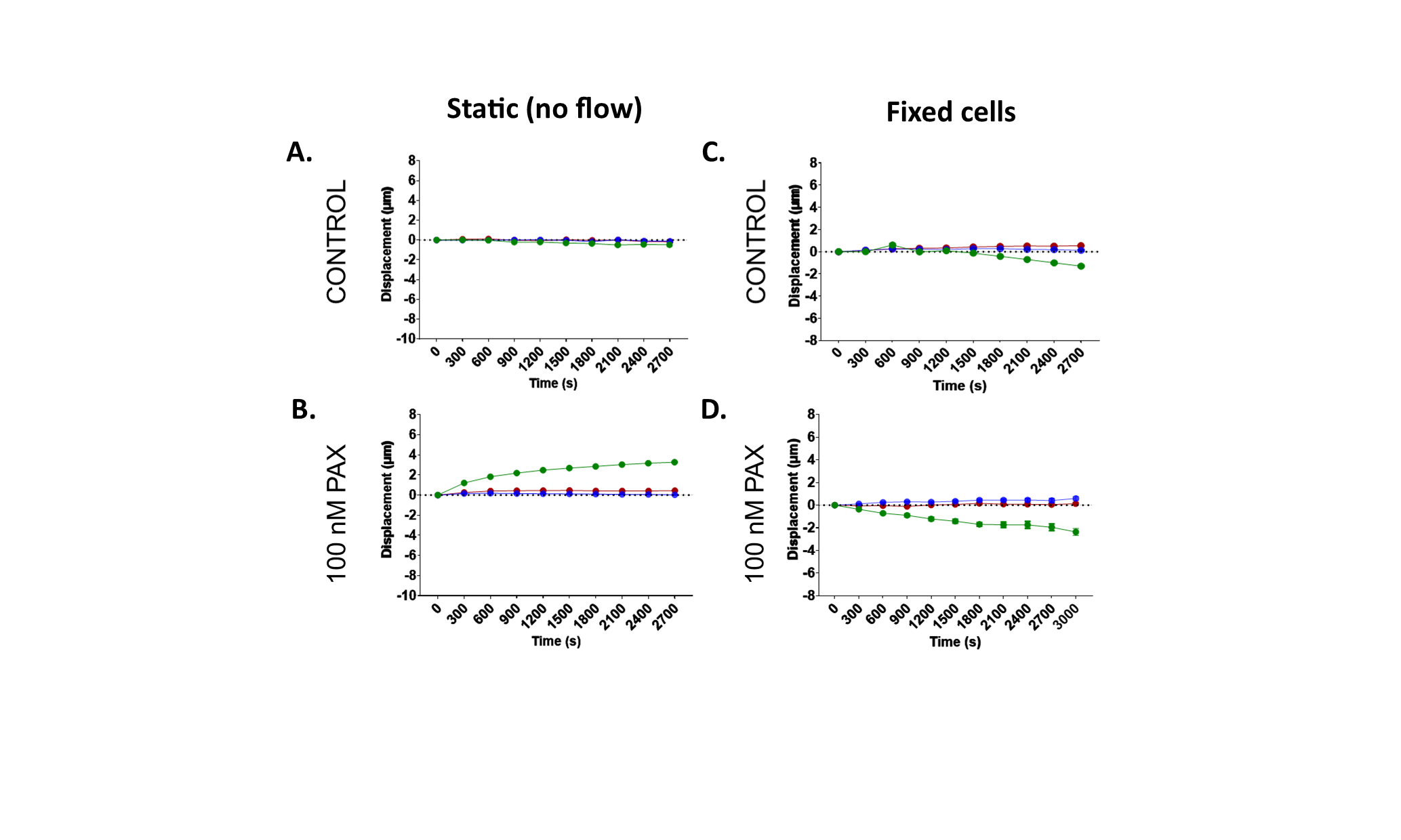

**Figure S1. Microbeads displacement profile in control (A) and microtubule-stabilized cells (B) under static condition.** When fixed cells were exposed to flow, displacement range remains small and similar to that of static (non-flow) condition for both control (C) and microtubule-stabilized cells (D). Red, blue and green lines represent the data in xyz.

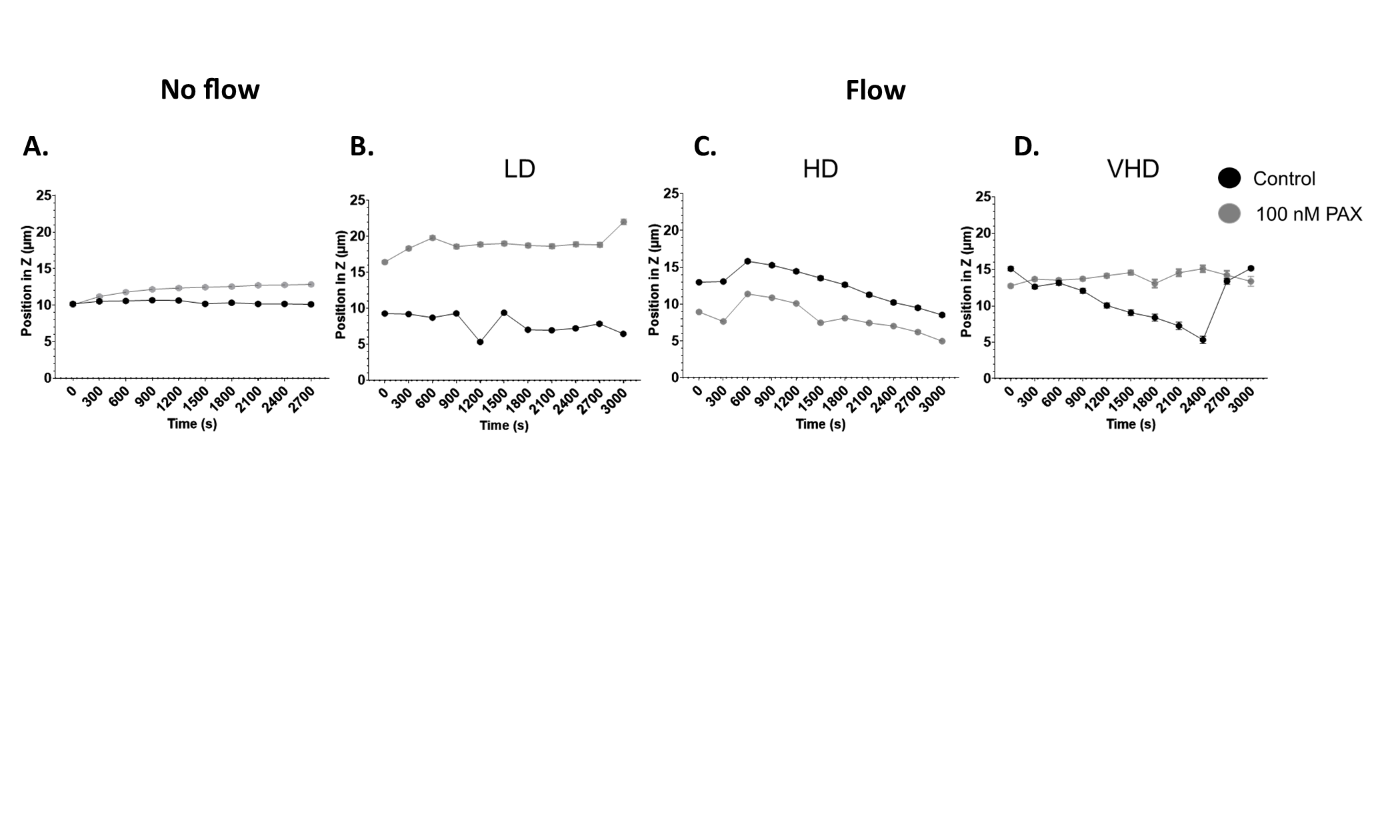

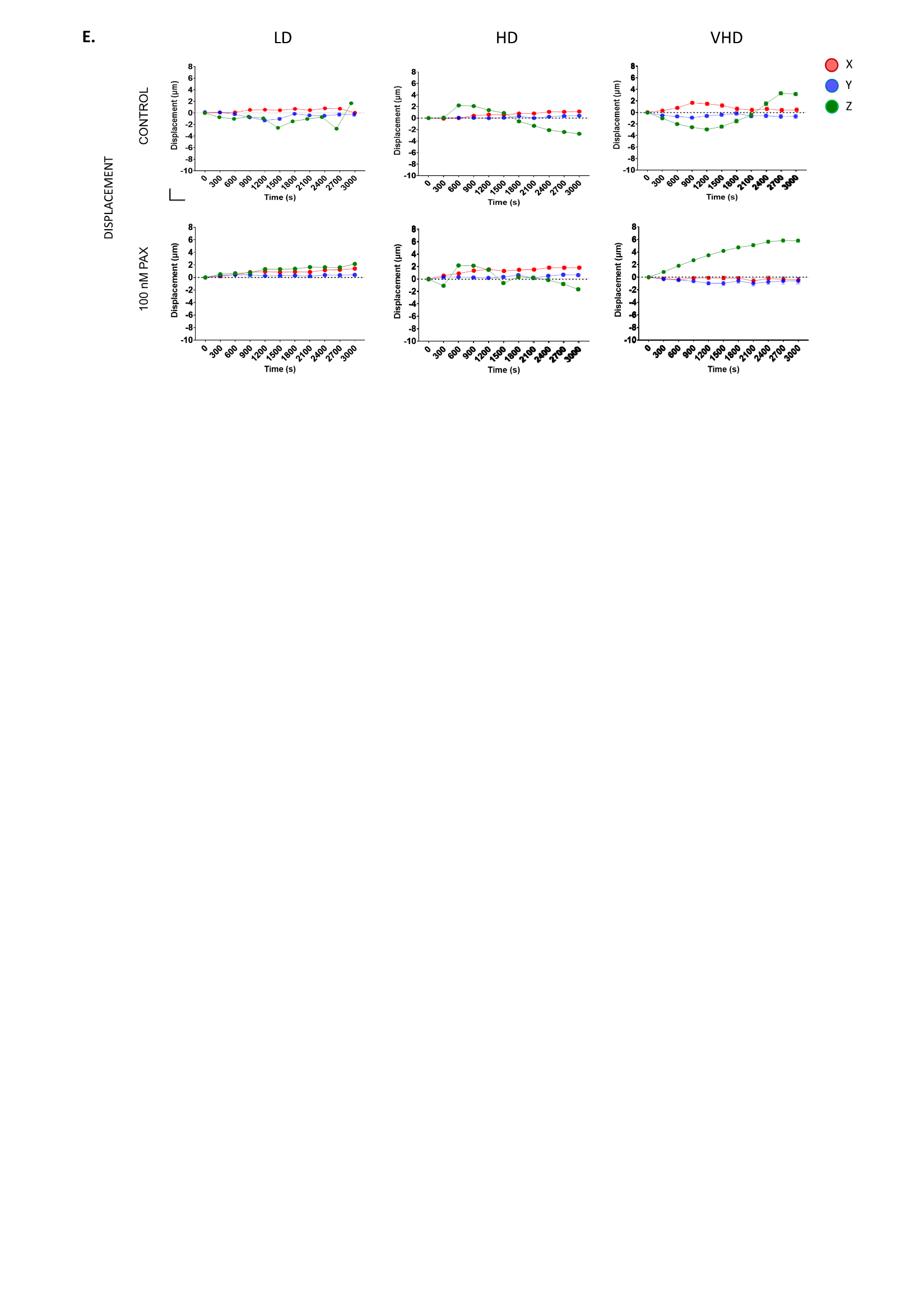

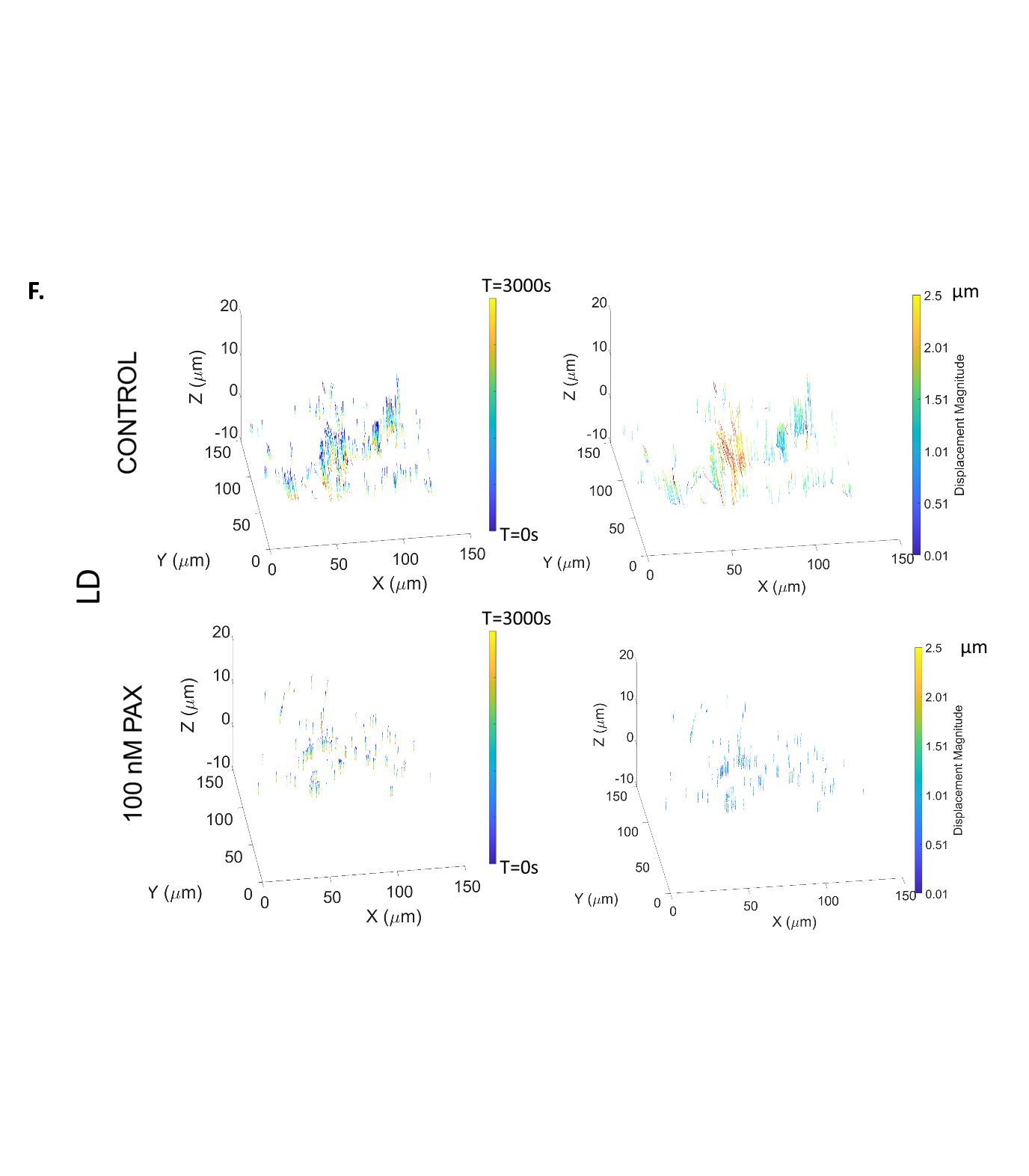

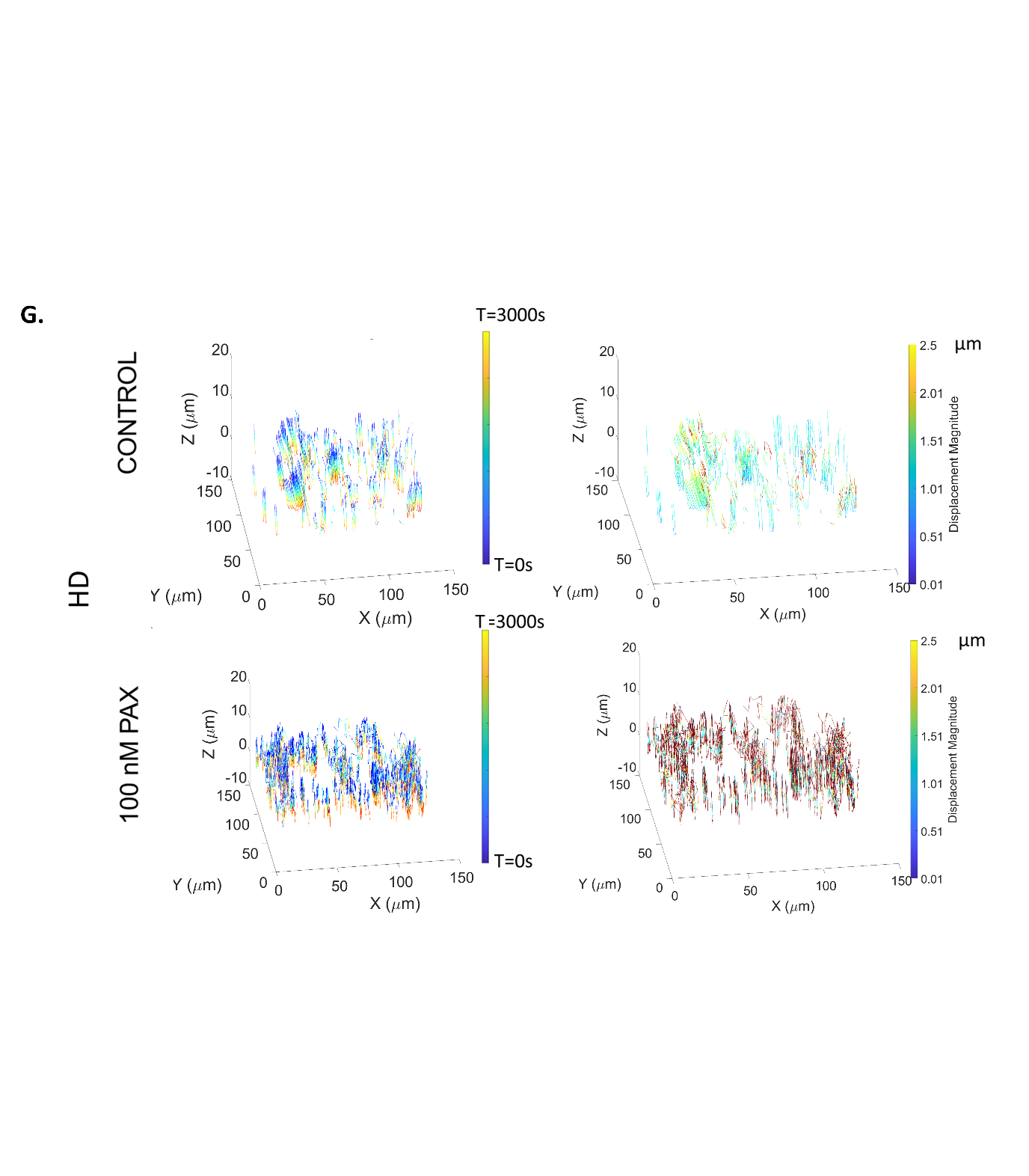

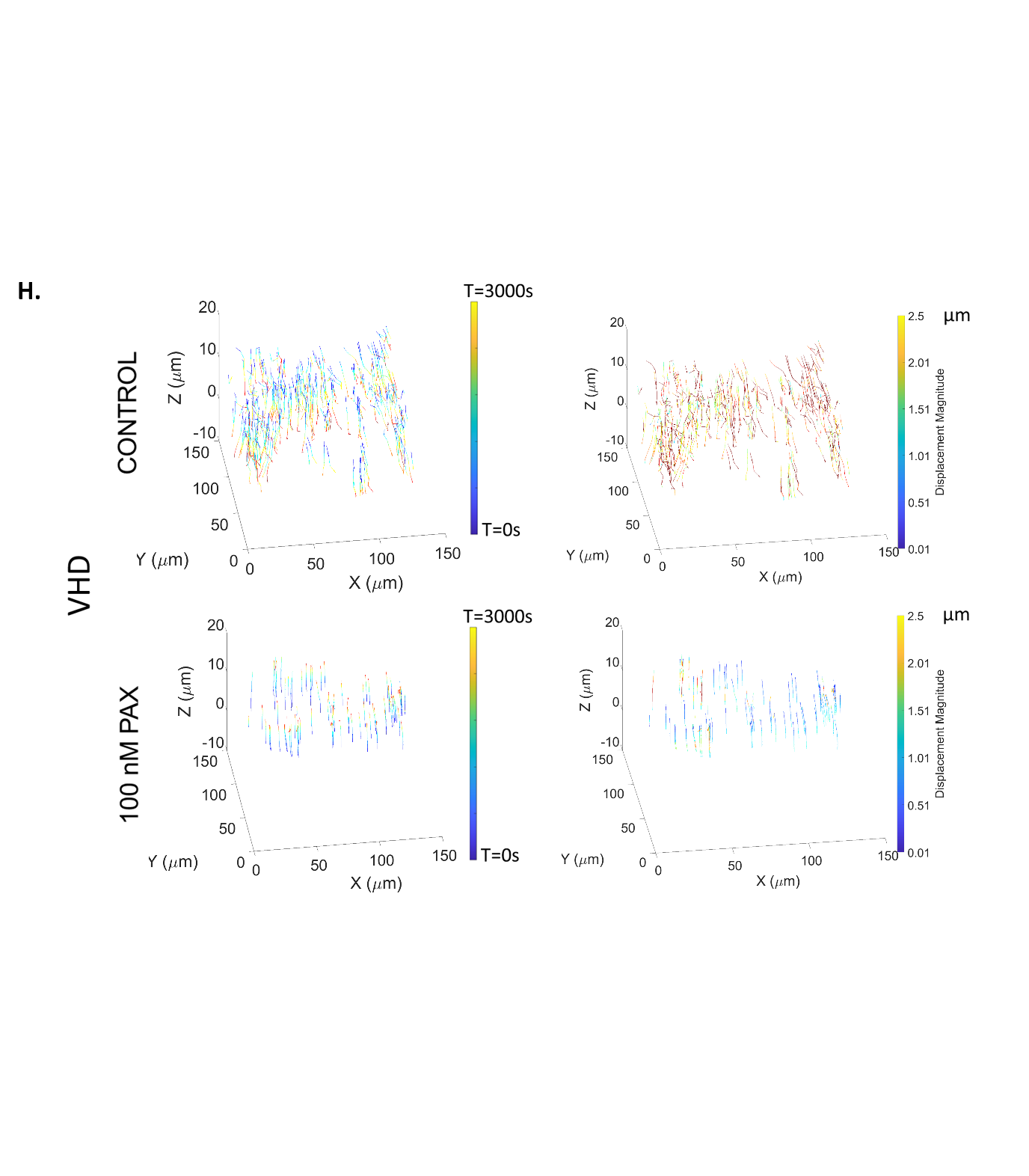

**Figure S2.** **Characterization of microbead displacement under flow.** (A) Cell height under static (non-flowed) condition is maintained over time. (B) Under flow at LD, cell height decreased over time more prominently in control cells than PAX-treated ones. At HD (C) and VHD (D), control cell height decreased more gradually than the PAX-treated cells. Changes in cell surface displacements occurred within an hour of flow and these changes relate to differences in cell seeding density as well as cell stiffness, shape and size (due to PAX treatment). Microbeads velocity in X, Y, Z was constant in PAX-treated over time but changes more dynamically in control cells. Microbeads velocity was slower after certain timepoints in LD cells both control and PAX treated. (E) Displacement profile of cell surface indicated the dynamic adaptation in cell position and movement between control and PAX-treated cells across the seeding density groups. Surface displacements of control cells were significantly more dynamic than that of the microtubule-stabilized cells for all the seeding density groups as shown in vector map of beads displacement. 3D Mapping of displacement vectors of cells tagged with fluorescent microbeads at (F) low density (LD), control cells exhibit larger and gradual displacement in Z than PAX-treated cells. (G) At high density (HD), PAX -treated cells displaced largely in Z and (H) very high density (VHD) cells fluctuated in position more prominently in PAX-treated cells. Left column shows displacement over time and right column shows displacement magnitude.

COMPUTATIONAL FLOW

**Validation of CFD with experimental data**

The ProFlow^TM^ chamber’s capacity to deliver highly controlled, laminar flow was tested previously by Anderson and Knothe Tate in published studies^11^. The analogous to the previous studies, fluid flow and delivery of controlled mechanical cues to cells was first simulated computationally within the flow chamber to predict the velocity needed to deliver controlled mechanical cues to the cells seeded within (Fig. S5.3A, B, C). The velocity and wall shear stress were calculated for three conditions of input pressure: 4, 20, and 40 Pa, to achieve increasing target shear stress (Supplementary Information).

Consistent with the previously reported performance of the flow chamber^11,14,17^ in the current 3D CFD analysis, we achieve constant velocity and shear stress within 78% and 58% of the middle volume of the chamber respectively. In the midplane across the x axis, the maximum shear stress is achieved within the 5 mm from inlet and the outlet and constant magnitude throughout the rest of the chamber (Fig. S5.3D, E, F). The maximum velocity across the x-axis is highest in the region closest to the inlet or outlet, and across the y-axis is highest within the middle region from few micrometres away from the wall. Across the height of the chamber (z-axis, 250 μm), the shear stress is minimum at the middle most z-height while velocity is highest at the furthest distance from the wall (Fig. S5.3G, H, I).

To validate the computational data of the flow chamber performance for the shear stress profile (Fig. S5.3J, K, L) and velocity (Fig S5.3. M, N, O) within the chamber, experimental data from the microbeads velocity was used as a reference. Cell culture medium containing microbeads was flowed through the empty flow chamber at a rate of 0.134 ml/min (previously predicted to induce shear stress of 0.2 dyn/cm^2^)^14^. Comparable to the model with 4 Pa input pressure at the inlet, the microbeads move at a mean velocity of 0.001131159 m/s in X direction (Fig. S5.3P), -0.000522999 m/s in Y (Fig. S5.3Q), and 0.002211235 m/s in Z (Fig. S5.3R).

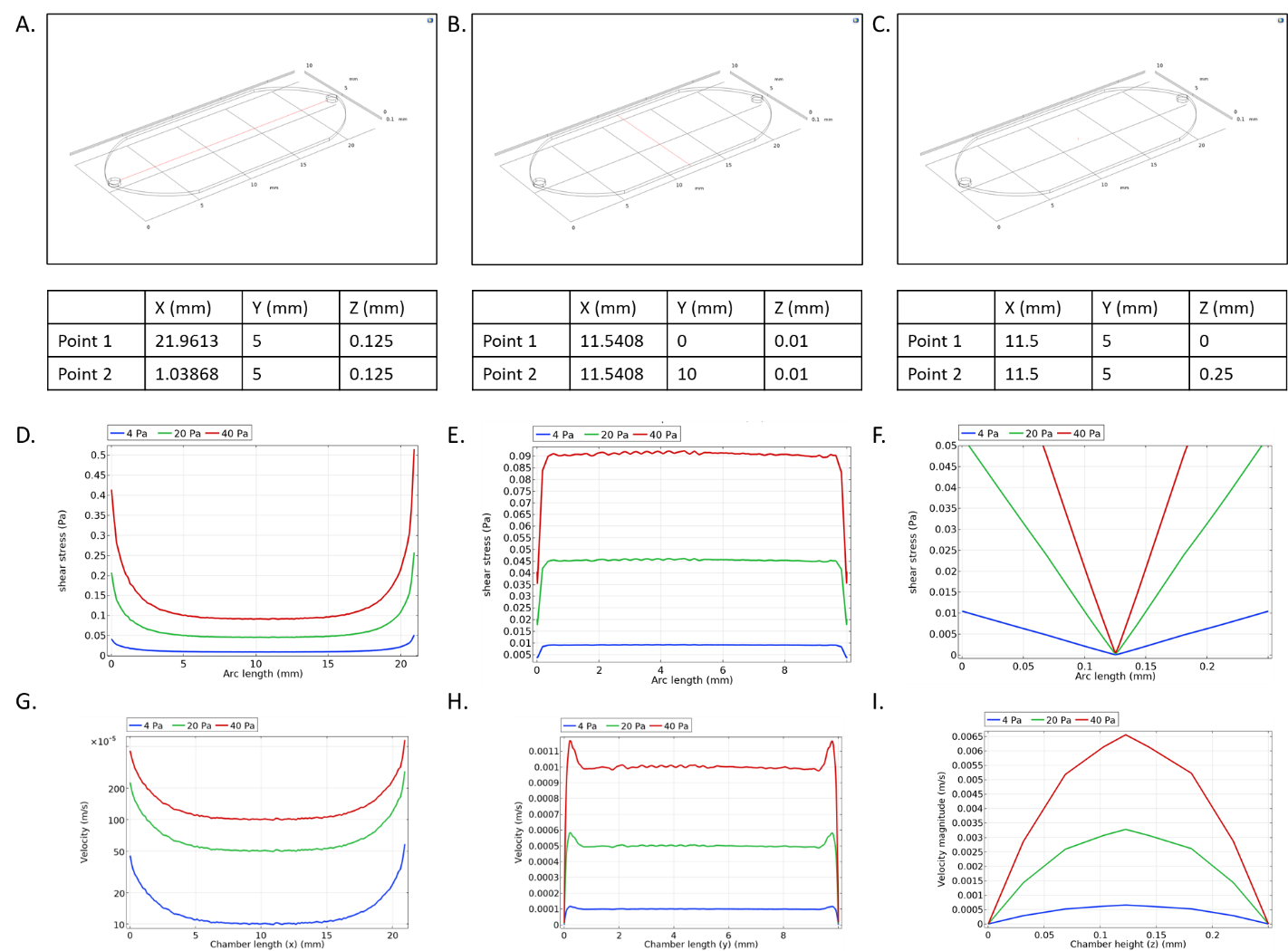

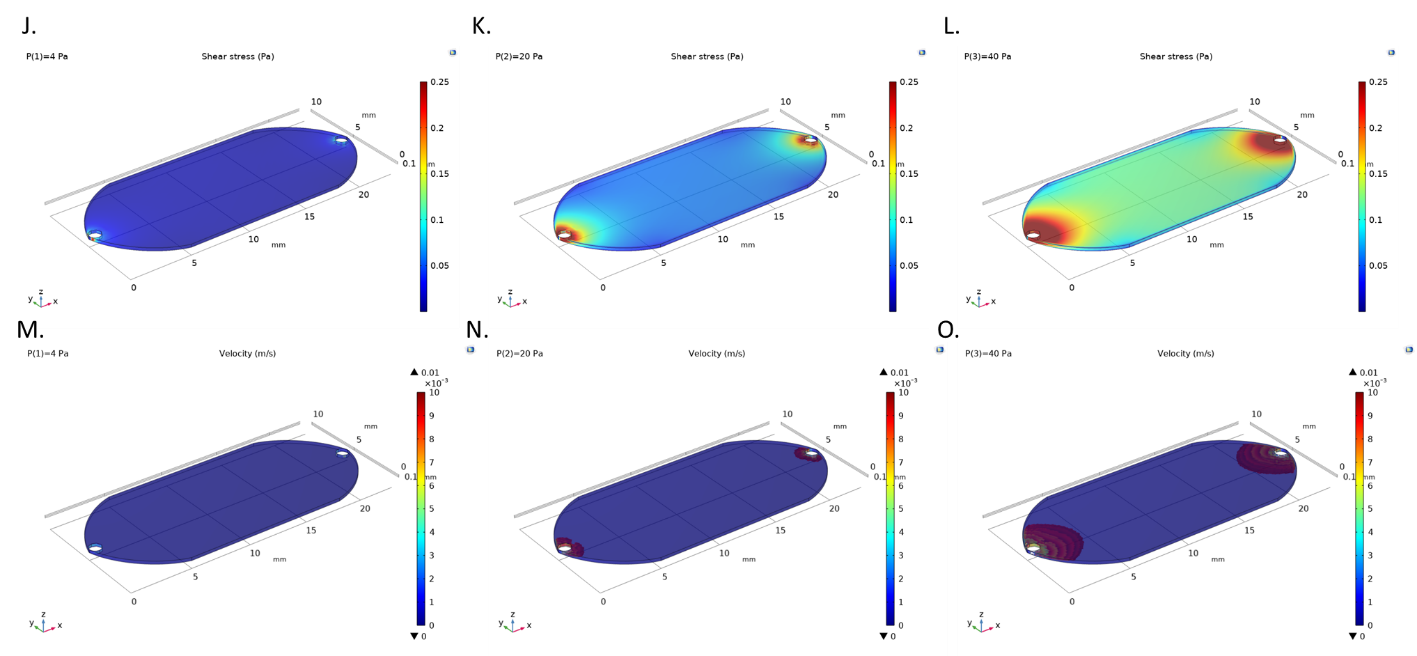

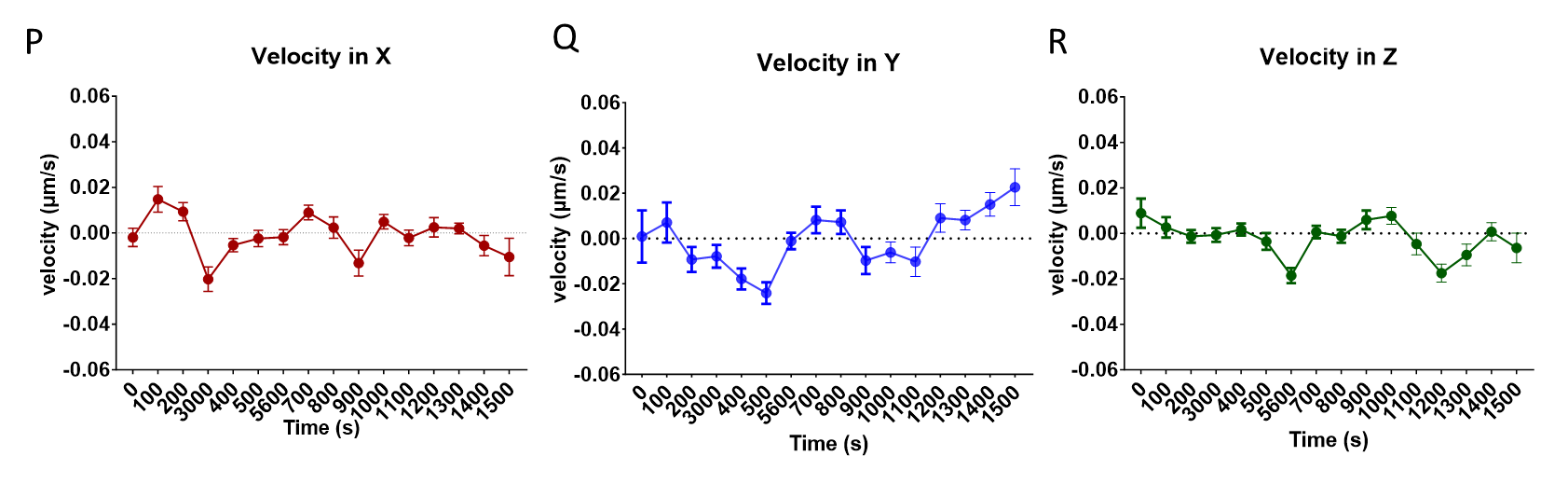
**Figure S3. Validation of the computational model of the ProFlow chamber previously reported in Anderson and Knothe Tate 2007**^11,23^ **and Song et al 2012**^14^**.** The eye-shaped flow chamber was recreated, and laminar flow was simulated by assigning sufficient input pressure at the inlet. The shear stress and velocity profile across the chamber (A) length (x axis), (B) width (y axis), and (C) height (z axis). Across the x-axis length at the midplane, within the middle part of the chamber, the shear stress is shown to be the lowest at the furthest from the inlet and outlet. Parametric sweep performed to introduce 4, 20, and 40 Pa input pressure at the inlet to initiate shear stress of 0.2, 1 and 2 dyn/cm^2^ of shear stress respectively, demonstrates the increase in shear stress magnitude that translates to five or ten times increase in stress. Flow moves from left (inlet) to right (outlet). The shear stress (D-F) and velocity (G-I) plotted along the chamber midplane in x and y (left and middle) and across the overall height in z (right). With increasing input pressure gradient, the middle 58% area of the chamber could deliver highly constant shear stress (J, K, L), and 70% of constant velocity (M, N, O). Flow velocity was validated experimentally by flowing fluorescent microbeads and tracking their position using time lapse imaging. Considering the random Z-position of microbeads during the flow, velocity within the chamber at a flow rate of 0.134 ml/min shows a comparable velocity profile in x (P), y (Q), and z (R), and to those modelled computationally.

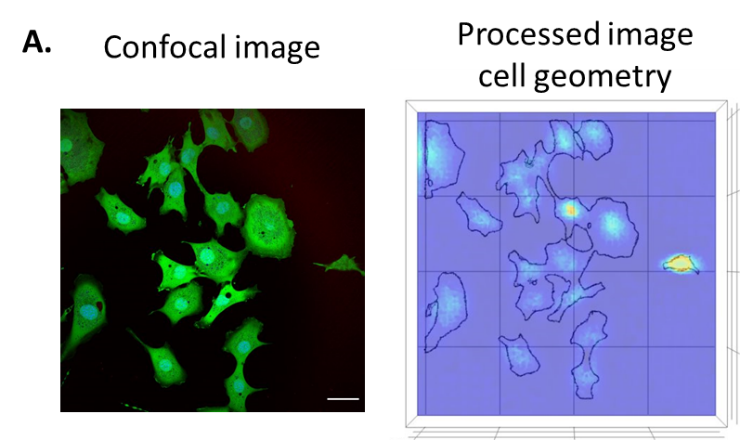

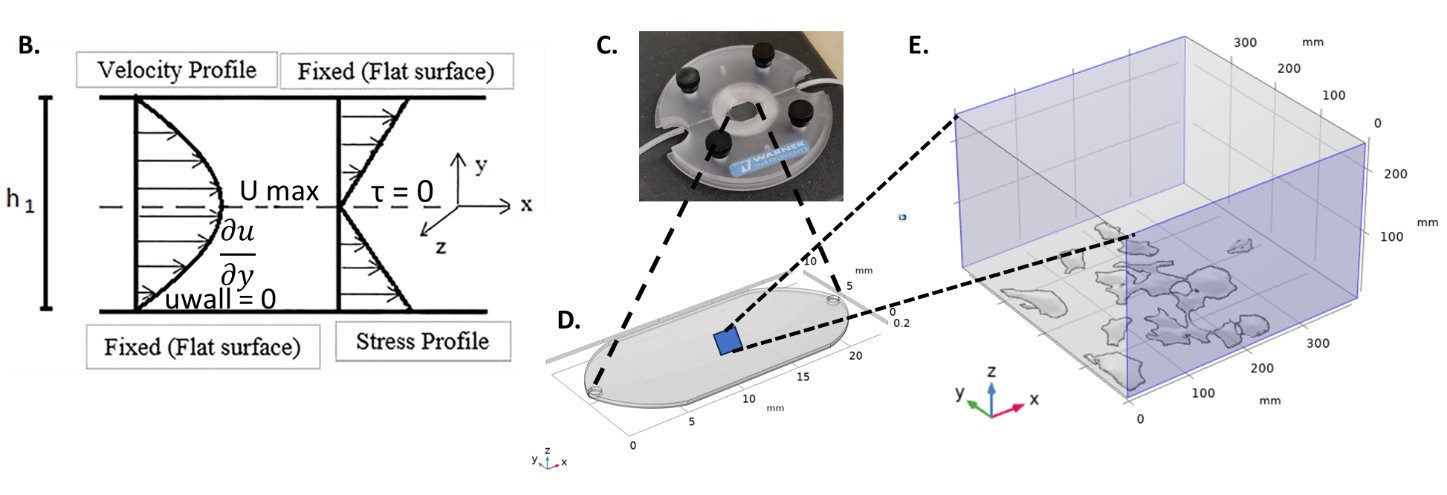

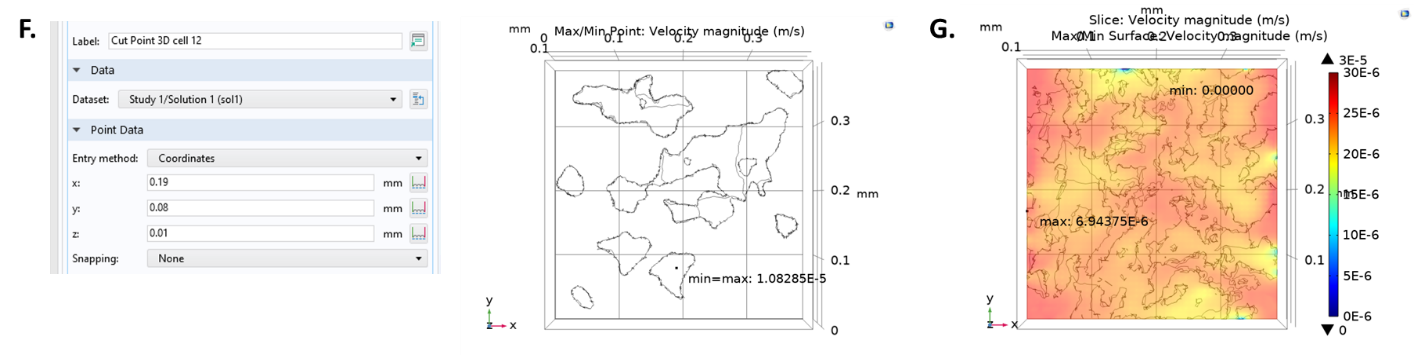

**Figure S4. Processing and importing cell structures and submodelling of ProFlow chamber**. (A) Processing of 3D confocal images of cells to build the geometry involves smoothing of cell surface, sacrificing small cell protrusions and irregular structures that would interfere with computational processes while preserving the cell structures. Scale bar = 50 µm. Schematic of the ProFlow chamber (B) with the eye-shaped gasket that defines the geometry of the chamber (C). The middle most region of the chamber is then sub-modelled to include 3D cell structures captured during live cell imaging (D). (E) The typical flow between two fixed parallel plates where velocity is maximum at the distance furthest from the plates (Umax) and zero at the plates (Uwall). Shear stress (τ) is maximum at the wall and zero at the mid-height of the chamber. In COMSOL, the velocity is calculated by the expression “spf.U” and the wall shear stress by the expression “spf.sr*spf.mu”, where spf.sr is the shear rate and spf.mu is the solution viscosity. (F) Representative of ‘Cut Point 3D’ coordinate located on the single cell where velocity and shear stress are calculated at a specific "cut through" plane. (G) Representative z-slice in 3D model at the height where velocity and shear stress are calculated.

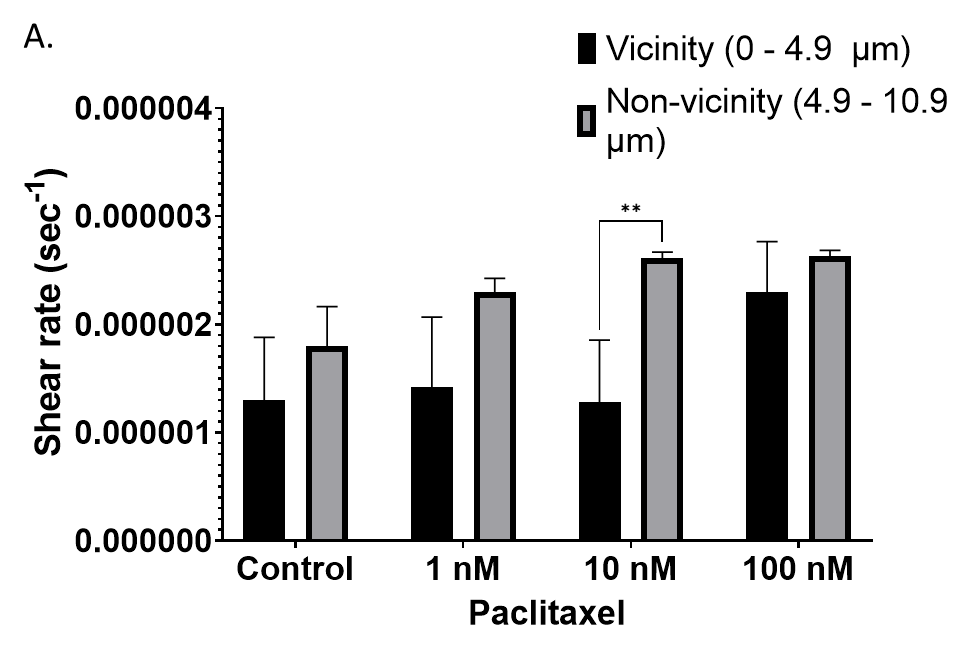

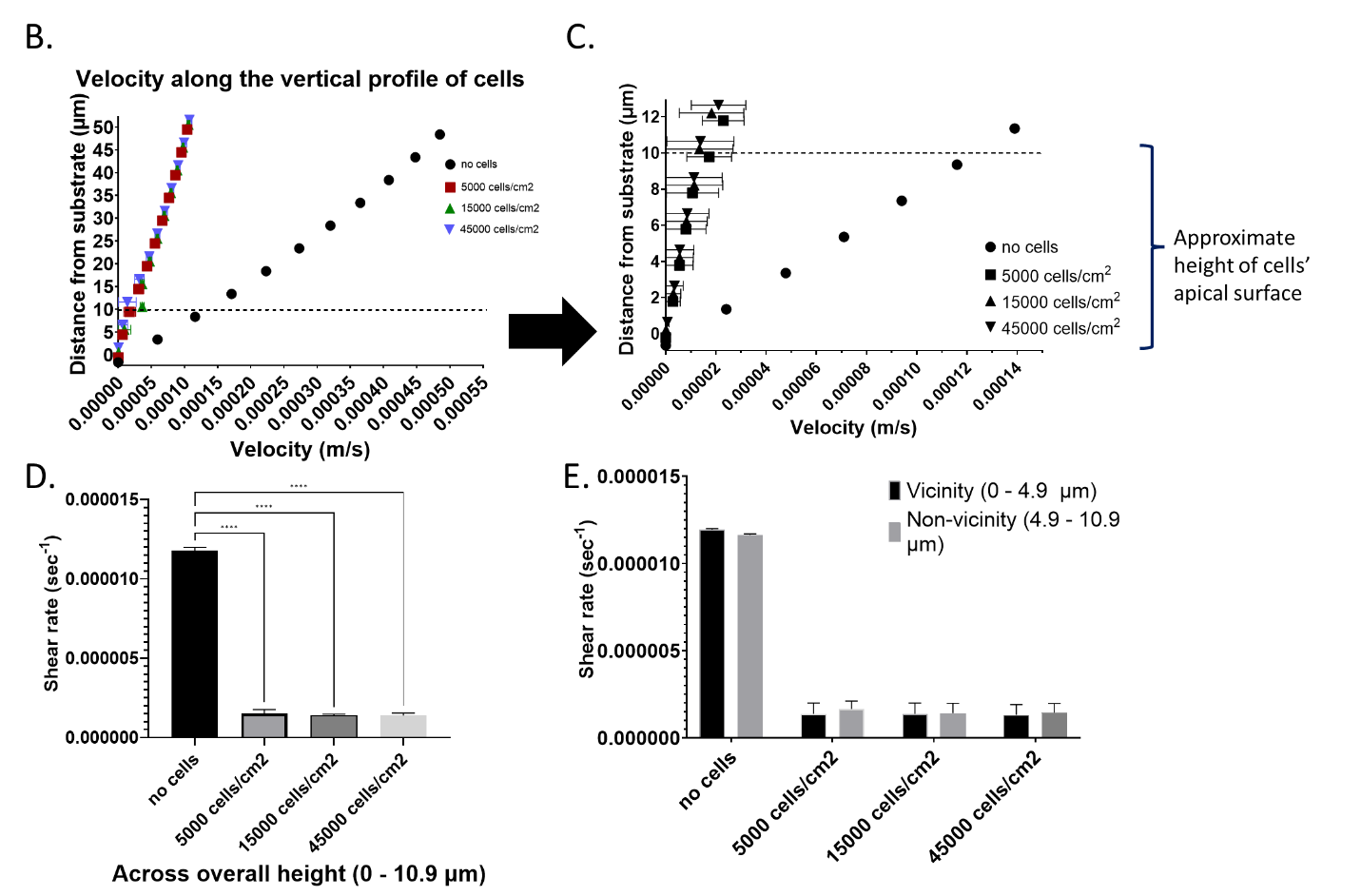

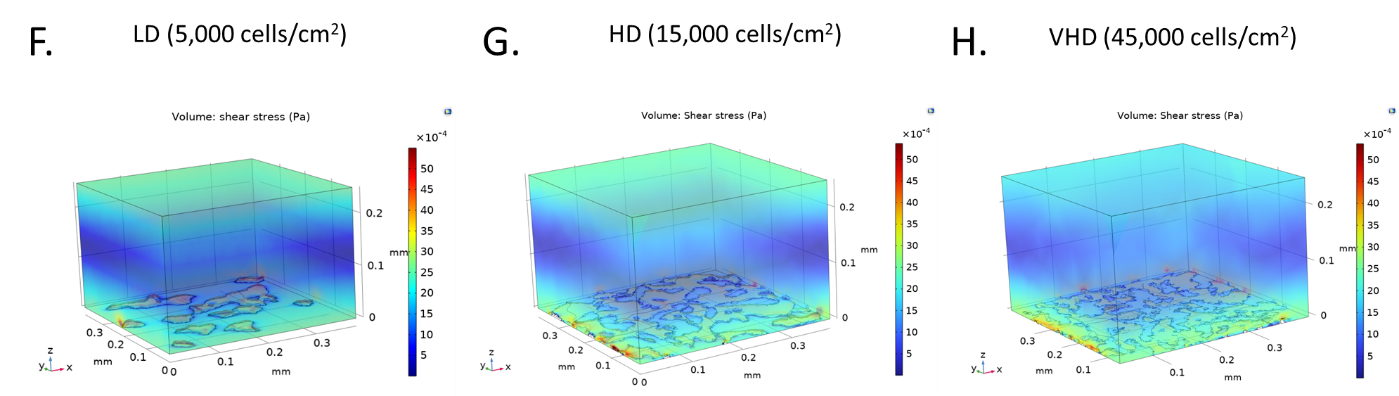

**Figure S5. Computational studies for mapping the flow fields around term cells treated with PAX or seeded at increasing seeding density.** (A) Shear rate increases along height (non-vicinity: apical and vicinity: basal) for those treated with low PAX concentration, but no significant difference in those of high PAX concentration. (B) Prediction of the velocity gradient and shear rate across the height of the chamber in the absence and presence of the cells with increasing seeding densities. Computational flow velocity along the one-fifth height of the chamber (C) and the vertical profile of the cells (0 – 10.9 um) (D). The shear rate varies significantly with respect to cell seeding density and cell height, and significant difference between the stresses at apical (non-vicinity) and basal (vicinity from substrate) region is observed in the low-density group (E). The shear stress profile varies significantly in relation to the presence of cells across the various seeding densities; LD (F) HD (G), and VHD (H) in the bottom of chamber.

SUPPLEMENTARY TABLES

**Supplementary Table S1. *Specification of material properties assigned in the model system and generated by COMSOL after computing for laminar flow physics.*** *Materials include cell culture medium (analogous to saline or water), cells (idealized as linear elastic material), and glass.*

| **Property** | **Variable** | **Expression** | **Unit** |
| --- | --- | --- | --- |
| **Fluid (cell culture medium idealized as saline)** | | | |
| Dynamic viscosity | μ | eta(T[1/K])[Pa*s] = 0.001 [Pa·s] | Pa·s |
| Density | rho | rho(T[1/K])[kg/m^3] = 1000[kg/m³] | kg/m³ |
| Ratio of specific heats | gamma | 1 | 1 |
| Electrical conductivity | sigma_iso_ ; sigma_ii_= sigmaiso, sigmaij = 0 | 5.5e-6[S/m] | S/m |
| Heat capacity at constant pressure | Cp | Cp(T[1/K])[J/(kg*K)] = 4.2 [J/(kg·K)] | J/(kg·K) |
| Thermal conductivity | k_iso ; kii = k_iso, kij = 0 | k(T[1/K])[W/(m*K)] = 0.6 [W/(m·K)] | W/(m·K) |
| **Cells** | | | |
| Density | rho | 1300 | kg/m³ |
| Dynamic viscosity | mu | 0.001 | Pa·s |
| Young's modulus | E | 2000 - 8000 | Pa |
| Poisson's ratio | nu | 0.36 | 1 |
| Elasticity matrix | {D11, D12, D22, D13, D23, D33, D14, D24, D34, D44, D15, D25, D35, D45, D55, D16, D26, D36, D46, D56, D66} ; Dij = Dji | {mat2.def.E*(1-mat2.def.nu)/((1+mat2.def.nu)*(1-2*mat2.def.nu)), mat2.def.E*mat2.def.nu/((1+mat2.def.nu)*(1-2*mat2.def.nu)), mat2.def.E*(1-mat2.def.nu)/((1+mat2.def.nu)*(1-2*mat2.def.nu)), mat2.def.E*mat2.def.nu/((1+mat2.def.nu)*(1-2*mat2.def.nu)), mat2.def.E*mat2.def.nu/((1+mat2.def.nu)*(1-2*mat2.def.nu)), mat2.def.E*(1-mat2.def.nu)/((1+mat2.def.nu)*(1-2*mat2.def.nu)), 0, 0, 0, 0.5*mat2.def.E/(1+mat2.def.nu), 0, 0, 0, 0, 0.5*mat2.def.E/(1+mat2.def.nu), 0, 0, 0, 0, 0, 0.5*mat2.def.E/(1+mat2.def.nu)} | Pa |
| Reference stress | sigRef | 1.2 | N/m² |
| Reference strain | eRef | 0.75 | 1 |
| Stress exponent | n_stress | 10^0.2 | 1 |
| **Silica glass** | | | |
| Relative permeability | mur_iso ; murii = mur_iso, murij = 0 | 1 | 1 |
| Electrical conductivity | sigma_iso ; sigmaii = sigma_iso, sigmaij = 0 | 1e-14[S/m] | S/m |
| Coefficient of thermal expansion | alpha_iso ; alphaii = alpha_iso, alphaij = 0 | 0.55e-6[1/K] | 1/K |
| Heat capacity at constant pressure | Cp | 703[J/(kg*K)] | J/(kg·K) |
| Relative permittivity | epsilonr_iso ; epsilonrii = epsilonr_iso, epsilonrij = 0 | 3.75 | 1 |
| Density | rho | 2203[kg/m^3] | kg/m³ |
| Thermal conductivity | k_iso ; kii = k_iso, kij = 0 | 1.38[W/(m*K)] | W/(m·K) |
| Young's modulus | E | 73.1e9[Pa] | Pa |
| Poisson's ratio | nu | 0.17 | 1 |
| Refractive index, real part | n_iso ; nii = n_iso, nij = 0 | 1.45 | 1 |
| Refractive index, imaginary part | ki_iso ; kiii = ki_iso, kiij = 0 | 0 | 1 |

***Supplementary Table S2. Parameter assignment for the laminar flow in the sub-model whereby values were derived from the full chamber model***

| **Name** | **Expression** | **Value** | **Description** |
| --- | --- | --- | --- |
| Re | 16 | 16 | Reynolds number |
| visc | 0.001[Pa*s] | 0.001 Pa·s | Fluid dynamic viscosity |
| dens | 1000[kg/m^3] | 1000 kg/m³ | Fluid density |
| U | 2.16667e-9[m^3/s] | 2.1667E-9 m³/s | Average inlet flow speed |
| H | 250[um] | 2.5E-4 m | Channel height |
| Re_check | dens*U*H/(visc) | 5.4167E-7 m² | Reynolds number |
| V | 0.00011[m/s] | 1E-4 m/s | Velocity (for fully developed flow) |
| sr | V/H | 0.4 1/s | shear rate |
| ss | sr*visc | 4E-4 Pa | target shear stress |
| P | 4[Pa] | 4 Pa | input pressure |
| ns | ss/H | 1.6 N/m³ | normal stress |

***Supplementary Table S3. Multivariate analysis and test between subject effects of the factorial interaction of all independent variables (seeding density, PAX concentration and flow) on the expression of actb and tuba1 gene****. Significance calculated at P<0.05 represent the significant dependence of actb and tuba1 expression on the independent variables or their interactions (highlighted in green).*

| **Tests of Between-Subjects Effects** | | | | | | |
| --- | --- | --- | --- | --- | --- | --- |
| Source |  | Type III Sum of Squares | df | Mean Square | F | **Sig.** |
| Corrected Model | actb | 467857.284^a^ | 11 | 42532.480 | 1.939 | **0.052** |
|  | tuba1 | 71975546.033^b^ | 11 | 6543231.458 | 1.775 | **0.079** |
| Intercept | actb | 5700102.521 | 1 | 5700102.521 | 259.844 | **0.000** |
|  | tuba1 | 61551902.748 | 1 | 61551902.748 | 16.698 | **0.000** |
| **seedingdens** | actb | 120151.954 | 2 | 60075.977 | 2.739 | **0.073** |
|  | tuba1 | 28665186.021 | 2 | 14332593.010 | 3.888 | **0.026** |
| **pax** | actb | 111298.751 | 1 | 111298.751 | 5.074 | **0.028** |
|  | tuba1 | 8949942.112 | 1 | 8949942.112 | 2.428 | **0.124** |
| **flow** | actb | 73428.907 | 1 | 73428.907 | 3.347 | **0.072** |
|  | tuba1 | 5173984.205 | 1 | 5173984.205 | 1.404 | **0.241** |
| **seedingdens * pax** | actb | 25758.871 | 2 | 12879.436 | 0.587 | **0.559** |
|  | tuba1 | 6226809.466 | 2 | 3113404.733 | 0.845 | **0.435** |
| **seedingdens * flow** | actb | 71339.664 | 2 | 35669.832 | 1.626 | **0.205** |
|  | tuba1 | 12824816.217 | 2 | 6412408.109 | 1.740 | **0.184** |
| **paxconc * flow** | actb | 64662.464 | 1 | 64662.464 | 2.948 | **0.091** |
|  | tuba1 | 2020166.870 | 1 | 2020166.870 | 0.548 | **0.462** |
| **seedingdens * pax * flow** | actb | 1216.673 | 2 | 608.336 | 0.028 | **0.973** |
|  | tuba1 | 8114641.142 | 2 | 4057320.571 | 1.101 | **0.339** |
| Error | actb | 1316195.894 | 60 | 21936.598 |  |  |
|  | tuba1 | 221174484.884 | 60 | 3686241.415 |  |  |
| Total | actb | 7484155.700 | 72 |  |  |  |
|  | tuba1 | 354701933.665 | 72 |  |  |  |
| Corrected Total | actb | 1784053.178 | 71 |  |  |  |
|  | tuba1 | 293150030.917 | 71 |  |  |  |
| a. R Squared = .262 (Adjusted R Squared = 0.127) | | | | | | |
| b. R Squared = .246 (Adjusted R Squared = 0.107) | | | | | | |

***Supplementary Table S4. Multivariate analysis and test between subject effects of the factorial interaction of all independent variables (seeding density, PAX concentration and flow) on the expression of gene markers of pre- (runx2, msx2) peri- (col1a1), and post-mesenchymal condensation (sox9, col2a1)****. Significance calculated at P<0.05 represent the significant dependence of gene expression on the independent variables or their interactions (highlighted in green).*

| **Tests of Between-Subjects Effects** | | | | | | |
| --- | --- | --- | --- | --- | --- | --- |
| Source |  | Type III Sum of Squares | df | Mean Square | F | **Sig.** |
| Corrected Model | runx2 | 34.723^a^ | 11 | 3.157 | 1.415 | **0.190** |
|  | col1a1 | 137477.523^b^ | 11 | 12497.957 | 2.791 | **0.00536369** |
|  | msx2 | .225^c^ | 11 | 0.020 | 2.120 | **0.03225837** |
|  | col2a1 | 1780.682^d^ | 11 | 161.880 | 3.551 | **0.00070649** |
|  | sox9 | 33688.255^e^ | 11 | 3062.569 | 3.857 | **0.00031699** |
| Intercept | runx2 | 394.898 | 1 | 394.898 | 177.009 | **0.00000000** |
|  | col1a1 | 1281287.681 | 1 | 1281287.681 | 286.082 | **0.00000000** |
|  | msx2 | 0.254 | 1 | 0.254 | 26.228 | **0.00000339** |
|  | col2a1 | 6233.478 | 1 | 6233.478 | 136.734 | **0.00000000** |
|  | sox9 | 91061.070 | 1 | 91061.070 | 114.692 | **0.00000000** |
| seedingdens | runx2 | 14.170 | 2 | 7.085 | 3.176 | **0.04886474** |
|  | col1a1 | 1444.233 | 2 | 722.116 | 0.161 | **0.85146196** |
|  | msx2 | 0.079 | 2 | 0.039 | 4.067 | **0.02206405** |
|  | col2a1 | 1083.393 | 2 | 541.696 | 11.882 | **0.00004495** |
|  | sox9 | 29354.051 | 2 | 14677.026 | 18.486 | **0.00000056** |
| pax | runx2 | 0.001 | 1 | 0.001 | 0.001 | **0.98067137** |
|  | col1a1 | 2324.535 | 1 | 2324.535 | 0.519 | **0.47405924** |
|  | msx2 | 0.034 | 1 | 0.034 | 3.555 | **0.06419755** |
|  | col2a1 | 5.422 | 1 | 5.422 | 0.119 | **0.73140709** |
|  | sox9 | 150.224 | 1 | 150.224 | 0.189 | **0.66513699** |
| flow | runx2 | 7.678 | 1 | 7.678 | 3.442 | **0.06848918** |
|  | col1a1 | 92901.179 | 1 | 92901.179 | 20.743 | **0.00002626** |
|  | msx2 | 0.033 | 1 | 0.033 | 3.434 | **0.06877138** |
|  | col2a1 | 23.217 | 1 | 23.217 | 0.509 | **0.47821991** |
|  | sox9 | 742.021 | 1 | 742.021 | 0.935 | **0.33755608** |
| seedingdens * pax | runx2 | 2.776 | 2 | 1.388 | 0.622 | **0.54026496** |
|  | col1a1 | 223.498 | 2 | 111.749 | 0.025 | **0.97536776** |
|  | msx2 | 0.026 | 2 | 0.013 | 1.340 | **0.26957569** |
|  | col2a1 | 89.858 | 2 | 44.929 | 0.986 | **0.37920160** |
|  | sox9 | 155.987 | 2 | 77.993 | 0.098 | **0.90658297** |
| seedingdens * flow | runx2 | 2.519 | 2 | 1.259 | 0.564 | **0.57163575** |
|  | col1a1 | 20242.461 | 2 | 10121.230 | 2.260 | **0.11318036** |
|  | msx2 | 0.038 | 2 | 0.019 | 1.963 | **0.14939758** |
|  | col2a1 | 42.712 | 2 | 21.356 | 0.468 | **0.62824150** |
|  | sox9 | 1112.614 | 2 | 556.307 | 0.701 | **0.50026587** |
| pax * flow | runx2 | 3.834 | 1 | 3.834 | 1.719 | **0.19487002** |
|  | col1a1 | 9820.050 | 1 | 9820.050 | 2.193 | **0.14390802** |
|  | msx2 | 0.006 | 1 | 0.006 | 0.664 | **0.41836531** |
|  | col2a1 | 167.174 | 1 | 167.174 | 3.667 | **0.06026983** |
|  | sox9 | 1781.620 | 1 | 1781.620 | 2.244 | **0.13937959** |
| seedingdens * pax * flow | runx2 | 3.745 | 2 | 1.873 | 0.839 | **0.43700647** |
|  | col1a1 | 10521.567 | 2 | 5260.783 | 1.175 | **0.31594053** |
|  | msx2 | 0.009 | 2 | 0.004 | 0.461 | **0.63279386** |
|  | col2a1 | 368.907 | 2 | 184.453 | 4.046 | **0.02247164** |
|  | sox9 | 391.738 | 2 | 195.869 | 0.247 | **0.78216481** |
| Error | runx2 | 133.857 | 60 | 2.231 |  |  |
|  | col1a1 | 268724.168 | 60 | 4478.736 |  |  |
|  | msx2 | 0.580 | 60 | 0.010 |  |  |
|  | col2a1 | 2735.309 | 60 | 45.588 |  |  |
|  | sox9 | 47637.665 | 60 | 793.961 |  |  |
| Total | runx2 | 563.479 | 72 |  |  |  |
|  | col1a1 | 1687489.373 | 72 |  |  |  |
|  | msx2 | 1.059 | 72 |  |  |  |
|  | col2a1 | 10749.469 | 72 |  |  |  |
|  | sox9 | 172386.990 | 72 |  |  |  |
| Corrected Total | runx2 | 168.580 | 71 |  |  |  |
|  | col1a1 | 406201.691 | 71 |  |  |  |
|  | msx2 | 0.805 | 71 |  |  |  |
|  | col2a1 | 4515.991 | 71 |  |  |  |
|  | sox9 | 81325.920 | 71 |  |  |  |
| a. R Squared = .206 (Adjusted R Squared = 0.060) | | | | | | |
| b. R Squared = .338 (Adjusted R Squared = 0.217) | | | | | | |
| c. R Squared = .280 (Adjusted R Squared = 0.148) | | | | | | |
| d. R Squared = .394 (Adjusted R Squared = 0.283) | | | | | | |
| e. R Squared = .414 (Adjusted R Squared = 0.307) | | | | | | |

***Supplementary Table S5. Multivariate analysis and test between subject effects of the factorial interaction of all independent variables (seeding density, PAX concentration and flow) on the expression of gene markers of angiogenesis (vegfa and pecam-1) and transcription factors for blood cell differentiation (cebpa)****. Significance calculated at P<0.05 represent the significant dependence of gene expression on the independent variables or their interactions (highlighted in green).*

| **Tests of Between-Subjects Effects** | | | | | | |
| --- | --- | --- | --- | --- | --- | --- |
| Source |  | Type III Sum of Squares | df | Mean Square | F | **Sig.** |
| Corrected Model | vegfa | 332.854^a^ | 11 | 30.259 | 4.594 | **0.00004860** |
|  | pecam1 | .003^b^ | 11 | 0.000 | 10.790 | **0.00000000** |
|  | cebpa | 18025.771^c^ | 11 | 1638.706 | 1.288 | **0.25379623** |
| Intercept | vegfa | 372.967 | 1 | 372.967 | 56.619 | **0.00000000** |
|  | pecam1 | 0.004 | 1 | 0.004 | 142.234 | **0.00000000** |
|  | cebpa | 6577.929 | 1 | 6577.929 | 5.168 | **0.02659576** |
| seedingdens | vegfa | 200.183 | 2 | 100.091 | 15.195 | **0.00000458** |
|  | pecam1 | 0.001 | 2 | 0.000 | 11.141 | **0.00007679** |
|  | cebpa | 11446.179 | 2 | 5723.089 | 4.497 | **0.01514707** |
| pax | vegfa | 0.013 | 1 | 0.013 | 0.002 | **0.96533253** |
|  | pecam1 | 0.001 | 1 | 0.001 | 54.458 | **0.00000000** |
|  | cebpa | 449.644 | 1 | 449.644 | 0.353 | **0.55449311** |
| flow | vegfa | 53.247 | 1 | 53.247 | 8.083 | **0.00609856** |
|  | pecam1 | 6.355E-05 | 1 | 6.355E-05 | 2.467 | **0.12152960** |
|  | cebpa | 883.722 | 1 | 883.722 | 0.694 | **0.40799572** |
| seedingdens * pax | vegfa | 14.028 | 2 | 7.014 | 1.065 | **0.35122372** |
|  | pecam1 | 0.001 | 2 | 0.000 | 10.283 | **0.00014454** |
|  | cebpa | 947.579 | 2 | 473.790 | 0.372 | **0.69075660** |
| seedingdens * flow | vegfa | 33.944 | 2 | 16.972 | 2.576 | **0.08443373** |
|  | pecam1 | 0.000 | 2 | 0.000 | 4.069 | **0.02201301** |
|  | cebpa | 1890.405 | 2 | 945.202 | 0.743 | **0.48017437** |
| pax * flow | vegfa | 7.149 | 1 | 7.149 | 1.085 | **0.30171759** |
|  | pecam1 | 0.000 | 1 | 0.000 | 3.916 | **0.05241404** |
|  | cebpa | 789.099 | 1 | 789.099 | 0.620 | **0.43414734** |
| seedingdens * pax * flow | vegfa | 24.290 | 2 | 12.145 | 1.844 | **0.16708074** |
|  | pecam1 | 0.000 | 2 | 8.840E-05 | 3.432 | **0.03880859** |
|  | cebpa | 1619.143 | 2 | 809.571 | 0.636 | **0.53289342** |
| Error | vegfa | 395.239 | 60 | 6.587 |  |  |
|  | pecam1 | 0.002 | 60 | 2.576E-05 |  |  |
|  | cebpa | 76364.649 | 60 | 1272.744 |  |  |
| Total | vegfa | 1101.060 | 72 |  |  |  |
|  | pecam1 | 0.008 | 72 |  |  |  |
|  | cebpa | 100968.349 | 72 |  |  |  |
| Corrected Total | vegfa | 728.093 | 71 |  |  |  |
|  | pecam1 | 0.005 | 71 |  |  |  |
|  | cebpa | 94390.420 | 71 |  |  |  |
| a. R Squared = .457 (Adjusted R Squared = 0.358) | | | | | | |
| b. R Squared = .664 (Adjusted R Squared = 0.603) | | | | | | |
| c. R Squared = .191 (Adjusted R Squared = 0.043) | | | | | | |

***Supplementary Table S6. Multivariate analysis and test between subject effects of the factorial interaction of all independent variables (seeding density, PAX concentration and flow) on the expression of gene markers of adipogenesis (pparg), chondrogenesis (acan) and osteogenesis (sp7).*** *Significance calculated at P<0.05 represent the significant dependence of gene expression on the independent variables or their interactions (highlighted in green).*

| **Tests of Between-Subjects Effects** | | | | | | |
| --- | --- | --- | --- | --- | --- | --- |
| Source |  | Type III Sum of Squares | df | Mean Square | F | **Sig.** |
| Corrected Model | pparg | 3.905^a^ | 11 | 0.355 | 2.412 | **0.0148111** |
|  | acan | .010^b^ | 11 | 0.001 | 1.713 | **0.0921358** |
|  | sp7 | 1.237^c^ | 11 | 0.112 | 6.225 | **0.0000010** |
| Intercept | pparg | 45.569 | 1 | 45.569 | 309.604 | **0.0000000** |
|  | acan | 0.030 | 1 | 0.030 | 58.068 | **0.0000000** |
|  | sp7 | 1.422 | 1 | 1.422 | 78.725 | **0.0000000** |
| seedingdens | pparg | 2.061 | 2 | 1.031 | 7.002 | **0.0018484** |
|  | acan | 0.002 | 2 | 0.001 | 1.724 | **0.1871382** |
|  | sp7 | 0.732 | 2 | 0.366 | 20.280 | **0.0000002** |
| pax | pparg | 0.690 | 1 | 0.690 | 4.690 | **0.0343285** |
|  | acan | 0.000 | 1 | 0.000 | 0.533 | **0.4683718** |
|  | sp7 | 0.026 | 1 | 0.026 | 1.418 | **0.2383478** |
| flow | pparg | 0.381 | 1 | 0.381 | 2.586 | **0.1130633** |
|  | acan | 0.002 | 1 | 0.002 | 4.580 | **0.0364250** |
|  | sp7 | 0.051 | 1 | 0.051 | 2.822 | **0.0981790** |
| seedingdens * pax | pparg | 0.377 | 2 | 0.188 | 1.279 | **0.2856673** |
|  | acan | 9.993E-05 | 2 | 4.997E-05 | 0.098 | **0.9066435** |
|  | sp7 | 0.056 | 2 | 0.028 | 1.540 | **0.2226724** |
| seedingdens * flow | pparg | 0.303 | 2 | 0.151 | 1.029 | **0.3635889** |
|  | acan | 0.004 | 2 | 0.002 | 4.297 | **0.0180321** |
|  | sp7 | 0.324 | 2 | 0.162 | 8.983 | **0.0003868** |
| pax * flow | pparg | 0.088 | 1 | 0.088 | 0.600 | **0.4417153** |
|  | acan | 0.000 | 1 | 0.000 | 0.228 | **0.6350259** |
|  | sp7 | 4.933E-05 | 1 | 4.933E-05 | 0.003 | **0.9584904** |
| seedingdens * pax * flow | pparg | 0.005 | 2 | 0.003 | 0.019 | **0.9816683** |
|  | acan | 0.001 | 2 | 0.000 | 0.634 | **0.5339734** |
|  | sp7 | 0.047 | 2 | 0.024 | 1.314 | **0.2762699** |
| Error | pparg | 8.831 | 60 | 0.147 |  |  |
|  | acan | 0.031 | 60 | 0.001 |  |  |
|  | sp7 | 1.084 | 60 | 0.018 |  |  |
| Total | pparg | 58.305 | 72 |  |  |  |
|  | acan | 0.070 | 72 |  |  |  |
|  | sp7 | 3.742 | 72 |  |  |  |
| Corrected Total | pparg | 12.736 | 71 |  |  |  |
|  | acan | 0.040 | 71 |  |  |  |
|  | sp7 | 2.320 | 71 |  |  |  |
| a. R Squared = 0.307 (Adjusted R Squared = 0.180) | | | | | | |
| b. R Squared = 0.239 (Adjusted R Squared = 0.100) | | | | | | |
| c. R Squared = 0.533 (Adjusted R Squared = 0.447) | | | | | | |
